# Protein Friction and Filament Bending Facilitate Contraction of Disordered Actomyosin Networks

**DOI:** 10.1101/2021.02.23.432588

**Authors:** Alexander K. Y. Tam, Alex Mogilner, Dietmar B. Oelz

## Abstract

We use mathematical modelling and computation to investigate how protein friction facilitates contraction of disordered actomyosin networks. We simulate two-dimensional networks using an agent-based model, consisting of a system of force-balance equations for myosin motor proteins and semi-flexible actin filaments. A major advantage of our approach is that it enables direct calculation of the network stress tensor, which provides a quantitative measure of contractility. Exploiting this, we use repeated simulations of disordered networks to confirm that both protein friction and actin filament bending are required for contraction. We then use simulations of elementary two-filament assemblies to show that filament bending flexibility can generate contraction on the microscopic scale. Finally, we show that actin filament turnover is necessary to sustain contraction and prevent pattern formation. Simulations with and without turnover also exhibit contractile pulses. However, these pulses are aperiodic, suggesting that periodic pulsation can only be achieved by additional regulatory mechanisms.

## 1 Introduction

The mechanics of actomyosin networks govern essential cellular processes, including muscle contraction [1], cell division [2], and cell motility [3]. Assemblies of actin and myosin exhibit diverse structural organisation. In muscles, actin filaments are aligned in parallel to form sarcomeres, in which myosin-II motor proteins generate force in accordance with the sliding filament theory [1]. Alternatively, actin filaments form a disordered two-dimensional meshwork in the cell cortex, located below the membrane of living cells. These filaments are cross-linked by myosin motors, which exert forces that give rise to cortical tension and flow [4]. This cortex deformation subsequently determines cellular morphology and locomotion. Understanding the mechanisms by which myosin motors generate local forces is challenging, and can be investigated using mathematical modelling and computation.

The sliding filament mechanism provides a starting point for investigating contraction in disordered networks. Myosin motors attached to pairs of parallel actin filaments can generate either contraction or expansion, depending on filament orientation. A motor protein bound to a pair of filaments with barbed ends facing outwards will generate local contraction, as shown in Figure 1.1a. Conversely, the filaments generate expansion if pointed ends face outwards (Figure 1.1b). However, this sliding filament mechanism alone cannot explain net contraction in disordered networks, in which filaments can cross at arbitrary angles and in either configuration with equal probability. In these networks, there must be additional symmetry-breaking mechanisms that favour contraction over expansion.

**Figure 1.1:**
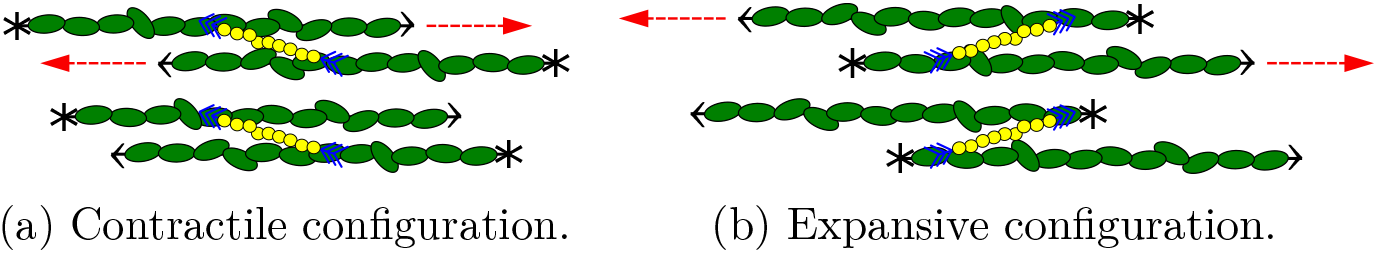
Schematic of the sliding filament mechanism for a myosin-II motor bound to a pair of actin filaments. Asterisks indicate filament barbed (plus) ends, arrow heads indicate pointed (minus) ends. Dashed arrows represent direction of filament movement.

**Figure 1.2:**
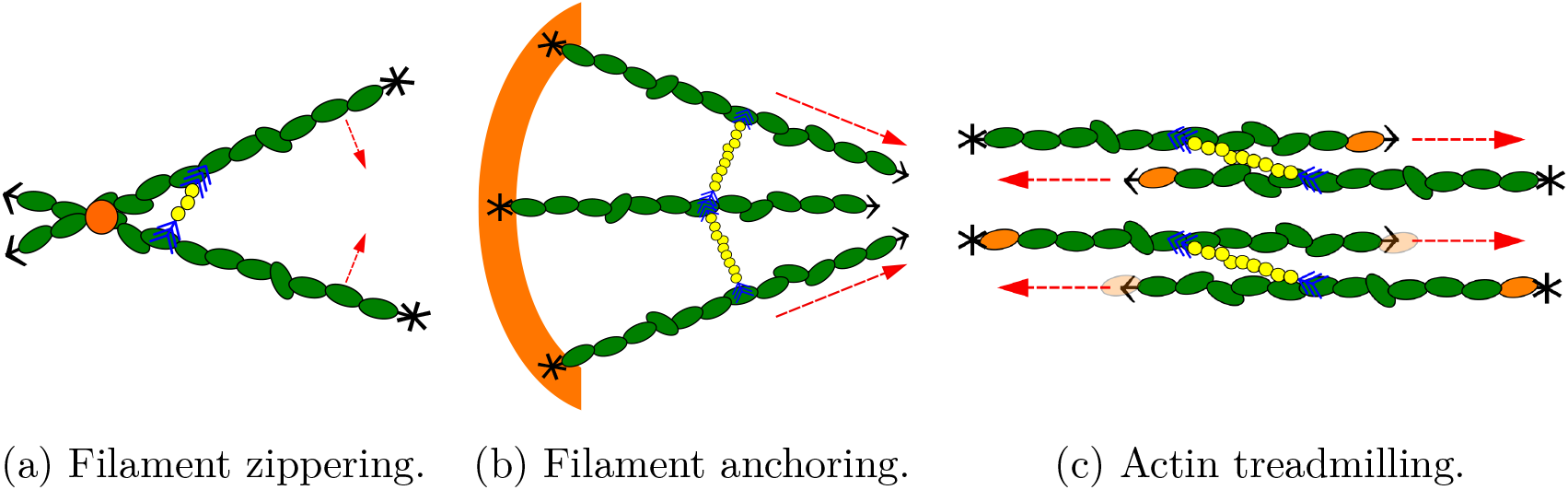
Schematic representations of proposed mechanisms for generating contraction via structural asymmetry.

Candidate mechanisms for generating contraction in disordered networks fall into the broad categories of structural and force asymmetries. Structural asymmetries break the random alignment of actin and myosin, enabling contractile configurations to emerge more often than expansive ones. Force asymmetries arise if filaments behave differently under tension and compression, enabling them to generate contraction more readily than expansion. In cells, mechanisms of contraction can be redundant and act as fail-safes in case network components are absent or lose function [5–7]. Many contractile mechanisms have been proposed and investigated for this reason.

Several hypotheses exist for generating structural asymmetries in two-dimensional networks. One example is a zippering mechanism, whereby a motor with non-zero length is displaced ahead of the intersection between two filaments (see Figure 1.2a). Motor movement towards the plus ends then pulls the filaments inwards, generating contraction [8–10]. Theoretical work by Lenz [9] showed that zippering can generate net contraction in disordered networks, but is unlikely to occur in practice. Another possible structural asymmetry is based on the observation that some filaments grow with barbed ends anchored to the cell membrane [5, 6] (see Figure 1.2b). Contraction can then occur via the sliding filament mechanism, since the anchored filaments are in a contractile alignment.

However, a drawback of this hypothesis is that only a small fraction of filaments in the cortex are anchored, such that non-anchored filaments are thought to play a major role in contractility [6]. A third hypothesised structural asymmetry for generating contraction is actin treadmilling, which involves simultaneous filament depolymerisation at minus ends and polymerisation at plus ends [11]. This enables contractile structures to persist as barbed ends are pulled inwards, generating a structural asymmetry (see Figure 1.2c). Oelz, Rubinstein, and Mogilner [12] showed that treadmilling generates network-scale contraction in one-dimensional ring-like geometry. Previous theoretical work has also shown that myosin motors lingering at filament barbed ends instead of unbinding can generate contraction [9, 13, 14]. However, although this behaviour has been observed in experiments, it is not known whether it occurs in non-muscle cells [15].

In contrast to these structural asymmetries, many studies consider a mechanism whereby filaments can sustain tension, but buckle under compression. The resulting asymmetric force propagation favours contraction. This has been illustrated *in vitro* [16] as well as theoretically [17] by suggesting that filaments nullify expansion by buckling when they are longer than a threshold length. Filament bending is likely to be relevant in cellular actomyosin, because the forces exerted by myosin motors are large enough to bend filaments with lengths below 1 μm [16], which is the approximate filament length [18]. However, the forces required to initiate bending are approximately 1000 times smaller than those required to rupture filaments [19]. Therefore, filament bending without buckling might also play a role in contraction.

Mathematical modelling has facilitated advancements in understanding actomyosin contraction. One phenomenological approach is to treat the actomyosin network as an active gel continuum [13, 20, 21]. In these models, filament and motor positions are expressed in terms of continuous density fields. Although these models can effectively predict pattern formation in actomyosin networks [22], many recent models focus on developing accurate microscopic descriptions of network components. Since we are interested in whether actin filament bending can induce contraction on both the microscopic and network scales, we focus on coarse-grained agent-based models. These models use simplified representations of individual network components, and track how they evolve over time. Agent-based models enable detailed description of the mechanics on a microscopic scale, and can subsequently be used to develop accurate continuum models [23].

Many agent-based models for the cytoskeleton exist, including publicly-available software Cytosim [24], AFINES [25], and MEDYAN [26]. These, and many other authors [7, 27–30], use modified Brownian dynamics to simulate actomyosin networks. Under this approach, actin filaments move according to an overdamped Langevin-like equation for the balance of forces between network components [7, 24–30]. Within this framework, many authors have recognised the importance of filament bending forces to contractility [7, 14, 25, 31–34]. A common approach is to focus on filament buckling [17, 25, 32, 35, 36] as a mechanism of contraction. This represents an extreme case of force asymmetry generated by deformable filaments. Using a one-filament worm-like chain model, Lenz [9] showed that motor-induced filament bending can generate contraction, and is relevant for typical experimental parameters. However, further quantitative analysis of this bending force asymmetry in filament networks is required.

Protein friction can be represented as effective viscous drag that acts point-wise at the binding site of a motor or cross-linker, or at the point of contact between filaments [37]. Using a one-dimensional model, Oelz, Rubinstein, and Mogilner [12] showed that a combination of actin treadmilling and drag distributed along filament pairs that overlap can contract a ring-like network of rigid filaments. In two-dimensions, protein friction manifests as point-wise drag at filament intersections [38, 39]. McFadden et al. [39] showed that point-wise drag and bending force asymmetry can generate contraction. These models with protein friction draw parallels between point-wise drag and cross-linkers [38, 39]. However, this implies that cross-linkers are either short and abundant, or turn over rapidly. The possibility of using point-wise drag to represent solid friction between filaments remains largely unexplored, and additional work is required to determine how this affects network contraction.

To address these research gaps, we develop a mathematical model for semi-flexible actin filaments and myosin motors to investigate how protein friction affects contractility. A promising simulation approach was developed by Dasanayake, Michalski, and Carlsson [40], and Hiraiwa and Salbreux [10], where the network configuration is given by the minimiser of a potential energy functional. However, these studies considered the evolution of random networks to a steady state, and neglected longer-time evolution of the network. In developing our model, we extend this approach to fully time-dependent simulations. An advantage of our approach is that it enables exact computation of the stress tensor components. This is in contrast to many previous models that use empirical methods to quantify contraction, for example observing filament velocities or summing force contributions of individual filaments.

## 2 Mathematical Model

We develop an agent-based model to simulate two-dimensional disordered networks. The network contains semi-flexible actin filaments, which we represent as finite-length curves in two-dimensional space. We represent myosin motors as dumbbells that behave as linear springs with equilibrium length zero, such that they attach to filament pairs at intersections. We assume that myosin motors detach immediately if they reach a filament plus end, and otherwise model force-dependent random detachment according to Bell’s law [41]. To maintain constant motor density, we do not model movement of unattached motors explicitly. Instead, we assume that a new motor immediately attaches at a random filament intersection when an unbinding event occurs. We then solve for the positions of filament nodes and myosin motors on a square domain with periodic boundary conditions to simulate network evolution.

Components in cytoskeletal networks undergo continuous turnover [31, 42–44]. This refers to the exchange of filaments, motors, and cross-linkers between the network and cytosol [10]. Turnover can occur when filament sever [16] or undergo treadmilling [12, 45, 46], which depend on motor [16] and cross-linker activity [35]. We explicitly model actin turnover by removing filaments (and any attached motors) at random with a constant rate [10, 39]. When a filament is removed, we immediately replace it with a new one at a random position, to maintain constant filament density. This represents a simple model for actin turnover, just as our treatment of myosin unbinding represents a simple model for motor turnover.

Protein friction is another mechanical feature that might influence network contractility [37, 47]. It can arise from binding and unbinding interactions between filaments and motors [47], filaments and cross-linkers [48], or from solid friction between filament pairs in contact [49]. Contact frictional forces are larger than hydrodynamic friction between filaments and the cytosol [48, 49], and have comparable magnitude to forces exerted by myosin motors [49]. In our model we apply viscous drag at intersections between actin filaments to model protein friction originating from either cross-linking or filament contact [38, 39]. We assume that presence of myosin motor prevents protein friction via cross-linking or filament contact, and do not apply point-wise drag between filament pairs connected to the same motor. Our model then enables investigation of whether protein friction, in conjunction with actin filament bending, can generate contraction.

We write the core model as a system of force-balance equations, which contains all mechanical features included in the model. In abstract terms, the system of equations is

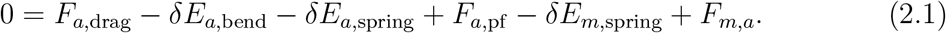

Actin filaments contribute to the force-balance via viscous drag, bending, stretching, and protein friction. Viscous friction penalises relative motion between actin filaments and the background medium, giving rise to drag forces *F*_*a*,drag_. We account for filament bending via the variation of *E*_*a*,bend_, which sums the elastic potential energy along the extent of each filament. The contribution of longitudinal spring forces, *E*_*a*,spring_, follows Hooke’s law with spring constant *k_a_*. Since actin filaments are effectively inextensible [50], we assume that *k_a_* is large. The symbol *F*_*a*,pf_ represents point-wise drag due to protein friction, which opposes relative motion of filament intersections.

The system (2.1) also contains two contributions relevant to myosin motors. Like for actin filaments, *E*_*m*,spring_ is the energy associated with longitudinal spring forces. These forces are governed by Hooke’s law with the spring constant *k_m_*, which we assume large to model the short length of myosin motors compared to actin filaments. For actin–myosin interactions we adopt a linear force–velocity relation for myosin motors, written as *F_m,a_*. Under this assumption, unloaded motors move at the velocity *V_m_*, and that motors cannot move if force exceeds the stall force, *F_s_*.

### 2.1 Numerical Method and Stress Calculation

In each simulation, we represent actin filaments as chains of six nodes, with adjacent nodes connected by straight line segments. We initialise filaments as straight entities with random centre positions and orientations, such that all nodes on the same filament are equidistant. Given the initial filament network, we place myosin motors at random intersections between filaments, such that each intersection accommodates a maximum of one motor. To evolve the network, at each time step we construct and minimise the energy functional,

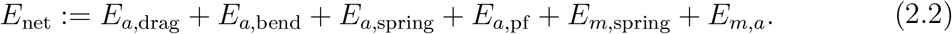

This functional includes pseudo-energy terms *E*_*a*,drag_, *E*_*a*,pf_, and *E_m,a_*, whose variations correspond to finite difference approximations of the force terms *F*_*a*,drag_, *F*_*a*,pf_, and *F_m,a_*, which cannot be interpreted as variations of potential energy. We then use the limited-memory Broyden–Fletcher–Goldfarb–Shanno (LBFGS) method to minimise (2.2) with respect to the positions of filament nodes and myosin motors. We perform this optimisation using the `Optim.jl` [51] package in Julia.

A key advantage of our method is that we compute the network forces exactly in numerical simulations. This differs from previous time-dependent Brownian dynamics models, in which contraction is often measured empirically, for example by observing filament velocities or by summing force contributions of individual filaments. We achieve this by adding extra terms to the energy functional, and defining the total energy

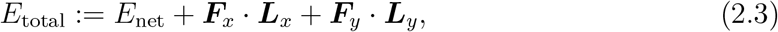

where ***L**_x_* = (*L_xx_,L_xy_*)^*T*^ and ***L**_y_* = (*L_yx_,L_yy_*)^*T*^ are vectors representing two edges of the domain. The vectors ***F**_x_* = (*F_xx_,F_xy_*)^*T*^ and ***F**_y_* = (*F_yx_,F_yy_*)^*T*^, illustrated in Figure 2.1, contain the normal and shear forces acting on the domain boundaries. In practice, we simulate the model on a domain of constant size. If we consider the vectors ***L**_x_* and ***L**_y_* as additional degrees of freedom, minimising (2.2) is equivalent to minimising (2.3), where the normal and shear force components are Lagrange multipliers that constrain the domain to constant size and shape. These forces quantify network behaviour, because they are equal and opposite to the forces generated in the network. In numerical simulations, we solve the model using (2.2), then compute ***F**_x_* = −δ_***L**_x_*_*E*_net_ and ***F**_y_* = −δ_***L**_x_*_*E*_net_ using Julia’s automatic differentiation capability (`AutoDiff.jl`). After calculating ***F**_x_* and ***F**_y_*, we combine the force components to compute the plane stress tensor,

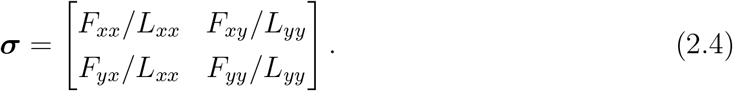

**Figure 2.1:**
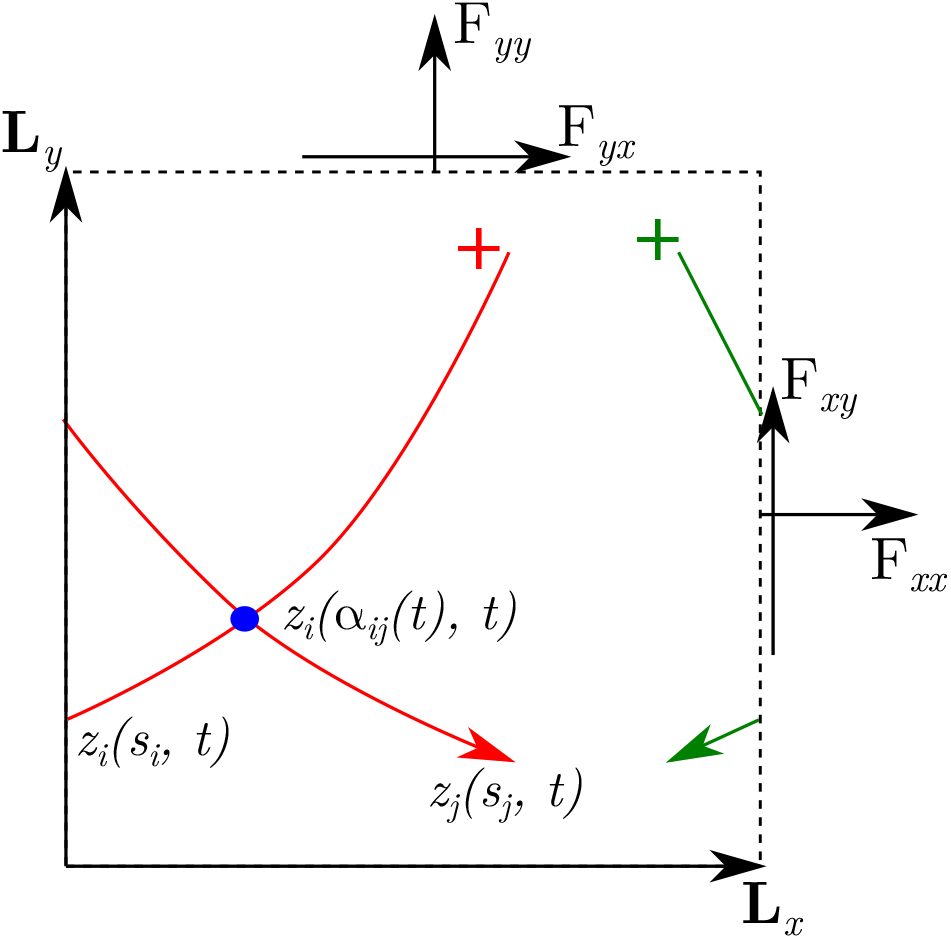
A schematic diagram of the periodic simulation domain, illustrating two actin filaments and a cross-linker. We use the domain vectors ***L**_x_*, and ***L**_y_*, and force components ***F**_x_*, and ***F**_y_*, to quantify network tension. The vectors 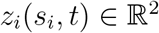 denote actin filament positions, parameterised by the arc length *s_i_*. The variable *α_ij_*(*t*) is the position of the intersection between the two filaments.

This fully describes the state of stress in the network at any time step. To obtain a single measure of contractility in a simulation, we define the time-averaged bulk stress

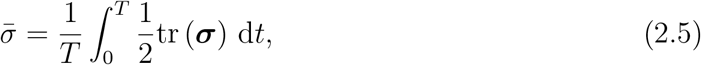

where *T* is the time over which the simulation runs, and tr(***σ***) is the trace of the stress tensor, which is invariant to co-ordinate rotations. The trace is also equal to the sum of the eigenvalues of ***σ***, and the associated eigenvectors indicate the principal stress directions. By convention, negative 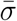 indicates contraction, and positive 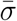 indicates expansion. Our method of quantifying network stress enables addition or removal of features from the energy functional, without changing the method of computing the forces. This flexibility is another advantage of our approach. Further details on the model derivation are provided in the supplementary section §A.

## 3 Results and Discussion

We use numerical simulations of our mathematical model to investigate how filament bending and protein friction affect contractility. In general, we simulate actomyosin networks using the default parameters provided in Table 3.1. Further details on the parameter estimation are provided in the supplementary section §A.3. We outline the main simulation results under subsequent headings.

**Table 3.1:**
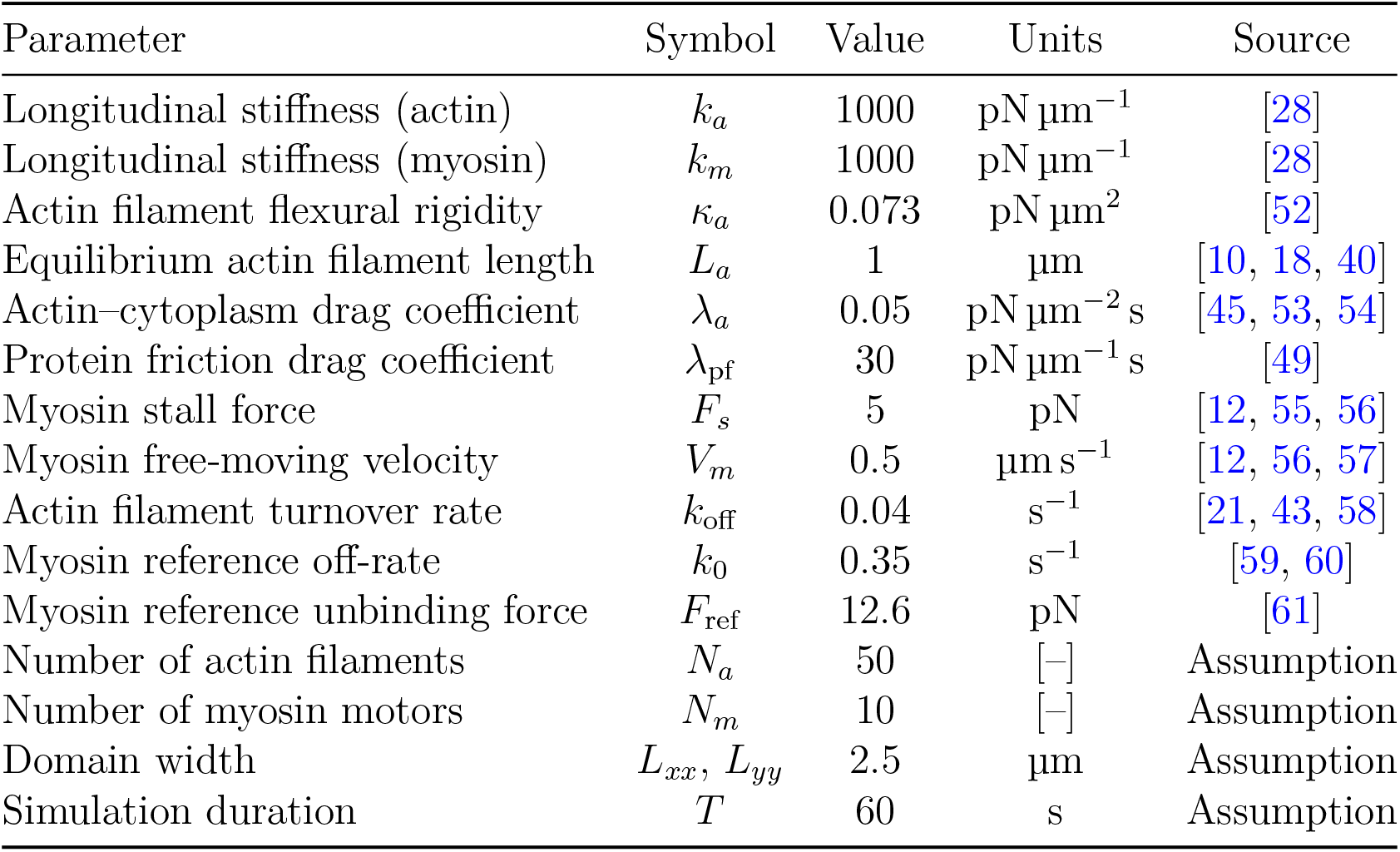
Default parameters for actomyosin network simulations.

| Parameter | Symbol | Value | Units | Source |
| --- | --- | --- | --- | --- |
| Longitudinal stiffness (actin) | $k_a$ | 1000 | $\text{pN } \mu\text{m}^{-1}$ | [28] |
| Longitudinal stiffness (myosin) | $k_m$ | 1000 | $\text{pN } \mu\text{m}^{-1}$ | [28] |
| Actin filament flexural rigidity | $\kappa_a$ | 0.073 | $\text{pN } \mu\text{m}^2$ | [52] |
| Equilibrium actin filament length | $L_a$ | 1 | $\mu\text{m}$ | [10, 18, 40] |
| Actin-cytoplasm drag coefficient | $\lambda_a$ | 0.05 | $\text{pN } \mu\text{m}^{-2} \text{ s}$ | [45, 53, 54] |
| Protein friction drag coefficient | $\lambda_{\text{pf}}$ | 30 | $\text{pN } \mu\text{m}^{-1} \text{ s}$ | [49] |
| Myosin stall force | $F_s$ | 5 | $\text{pN}$ | [12, 55, 56] |
| Myosin free-moving velocity | $V_m$ | 0.5 | $\mu\text{m s}^{-1}$ | [12, 56, 57] |
| Actin filament turnover rate | $k_{\text{off}}$ | 0.04 | $\text{s}^{-1}$ | [21, 43, 58] |
| Myosin reference off-rate | $k_0$ | 0.35 | $\text{s}^{-1}$ | [59, 60] |
| Myosin reference unbinding force | $F_{\text{ref}}$ | 12.6 | $\text{pN}$ | [61] |
| Number of actin filaments | $N_a$ | 50 | [-] | Assumption |
| Number of myosin motors | $N_m$ | 10 | [-] | Assumption |
| Domain width | $L_{xx}, L_{yy}$ | 2.5 | $\mu\text{m}$ | Assumption |
| Simulation duration | $T$ | 60 | $\text{s}$ | Assumption |

### 3.1 Actin Filament Bending Generates Bias to Contraction

To investigate actin filament bending as a contractile mechanism, we compared ten simulations of semi-flexible filaments with ten simulations of rigid filaments (neglecting bending energy), with all parameters as in Table 3.1. We then compared the time-averaged bulk stress (2.5) for *T* = 60s. These simulations reveal that bending is essential to generating contraction. As Figure 3.1 shows, the network contracted in each simulation with semi-flexible filaments, but always expanded with rigid filaments. The mean time-averaged bulk stresses across the simulations of 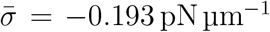 for semi-flexible filaments and 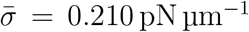 for rigid filaments confirms that bending generates systematic bias towards contraction.

**Figure 3.1:**
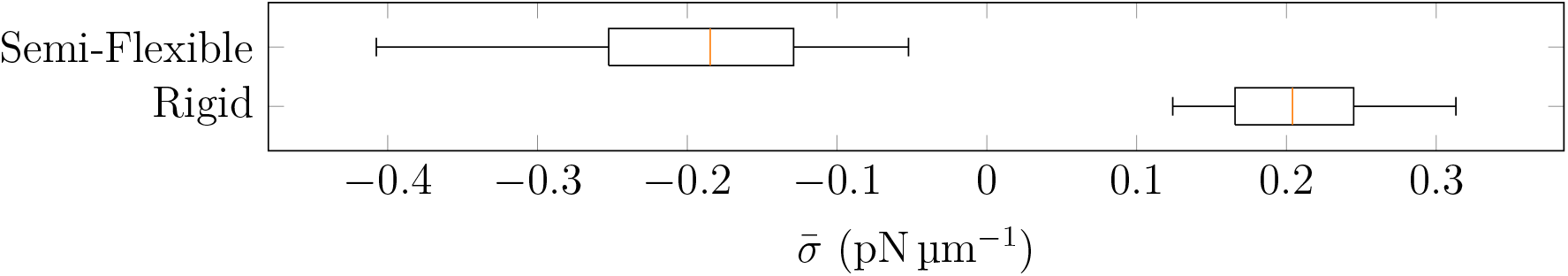
Box plots of time-averaged bulk stress 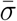 in ten semi-flexible simulations and ten rigid simulations, both using default model parameters. Box plots represent ten random simulations for a given parameter, and orange bars indicate median values.

We hypothesise that the magnitude of contraction depends on the extent of filament bending in the network. To investigate this, at each time step in the simulations we compute the local curvature

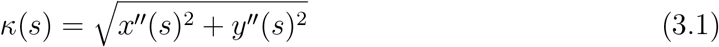

at each filament node. To obtain a measure of total curvature for one filament, we use the trapezoidal rule to integrate the curvature along the filament. We then obtain a single measure to characterise the extent of filament curvature by averaging integrated curvature over all filaments and time, defining

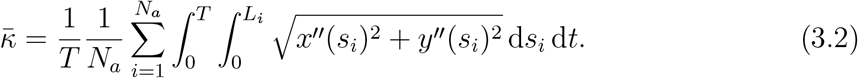

In the remainder of this manuscript, bar notation will represent quantities similarly averaged over filaments and time.

Since the extent of filament bending depends on the actin flexural rigidity, we varied *κ_a_* and investigated its effect on stress production. For each value of *κ_a_* tested, we ran ten random simulations and computed 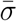. Box plots of network bulk stress presented in Figure 3.2a show that decreasing *κ_a_* increases contractility. This is expected, because *κ_a_* describes the resistance of filaments to bending. As Figure 3.2b shows, the increase in contractile stress that occurs with decreasing *κ_a_* corresponds to increased filament curvature. This accords with the hypothesis that filament bending generates force asymmetry, and subsequently contraction. Furthermore, the flexural rigidity for actin filaments, *κ_a_* = 0.073 pN μm^2^ [52], lies within the region for which we expect contraction. Actin filament bending is thus a plausible mechanism of contraction in biological cells.

**Figure 3.2:**
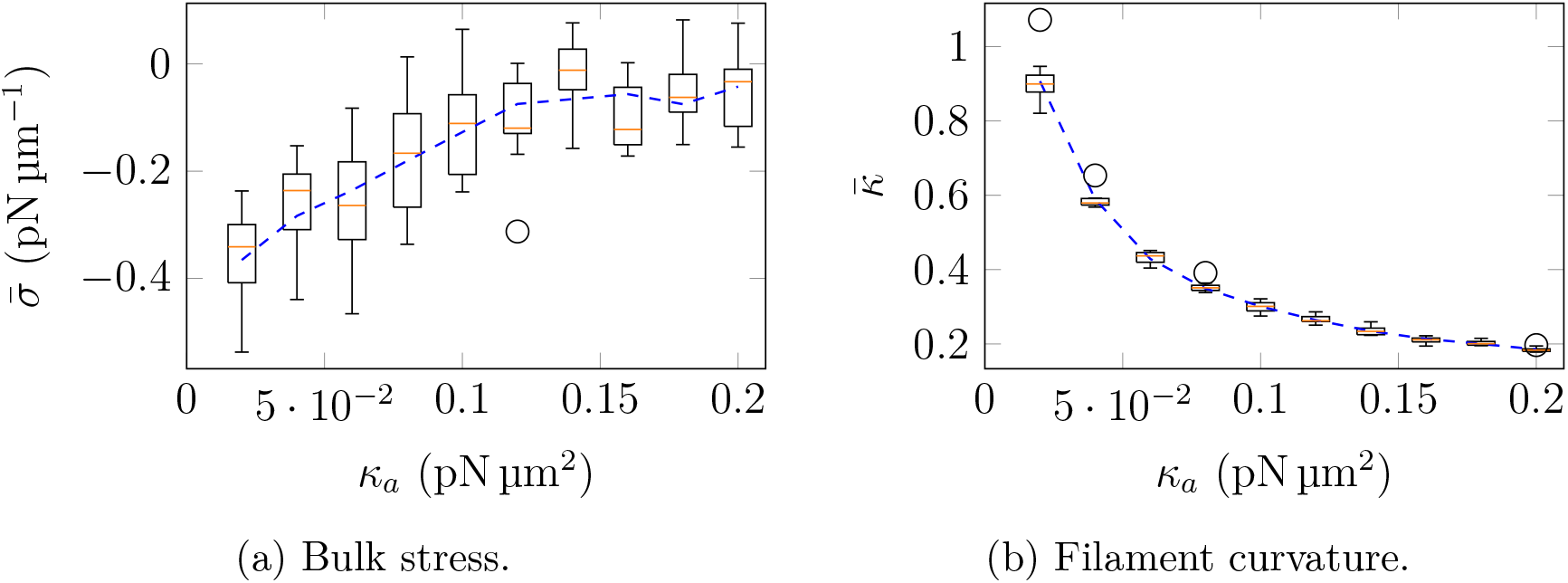
Simulation results for the effect of actin flexural rigidity, *κ_a_*, on time-averaged quantities of interest. Box plots represent ten random simulations for a given parameter, orange bars indicate median values, and dashed curve is the mean data, smoothed with a Savitsky–Golay filter.

### 3.2 Bending Generates Contraction on the Microscopic Scale

To better understand the microscopic mechanisms of contraction, we simulate two actin filaments with an attached myosin motor. Our objective is to determine whether filament bending generates force asymmetry in this simple structure, or whether contraction relies on network-scale interactions. In two-filament simulations, we use parameters in Table 3.1, except for *L_xx_ = L_yy_* = 2 μm, and *λ_a_* = 10 pN μm^−2^ s. A justification for the larger value of *λ_a_* is that we assume the two-filament structure is embedded in a dense, homogeneous background network. When a single fibre is immersed in such a network, protein friction manifests itself as drag acting uniformly along the entire filament length. The larger value of *λ_a_* then replaces protein friction at filament intersections, which cannot occur in the two-filament simulations because the motor occupies the only intersection.

We initialise the two filaments in a square domain, and characterise their positions by the angle *θ* ∈ [0,*π*], which is the angle between the two filaments measured at their intersection point. The relative motor positions are denoted by *m_1_* and *m_2_*, such that *m_i_* ∈ [0, *L_i_*] for *i* = 1, 2, measures the distance of the motor binding site from the minus end of filament *i*. Figure 3.3 illustrates this situation. We hypothesise that the extent of expansion or contraction of the two-filament structure depends on *θ*, *m*_1_, and *m*_2_. As the motor slides the filaments, it pulls filament branches between the motor and plus-ends together, generating contraction. Simultaneously, it pushes filament branches between the motor and minus-ends apart, generating expansion. Furthermore, the filaments will move the most if they are anti-parallel, or *θ = π*. Conversely, when filaments are parallel (*θ* = 0), the motor will traverse the filaments without generating relative motion.

**Figure 3.3:**
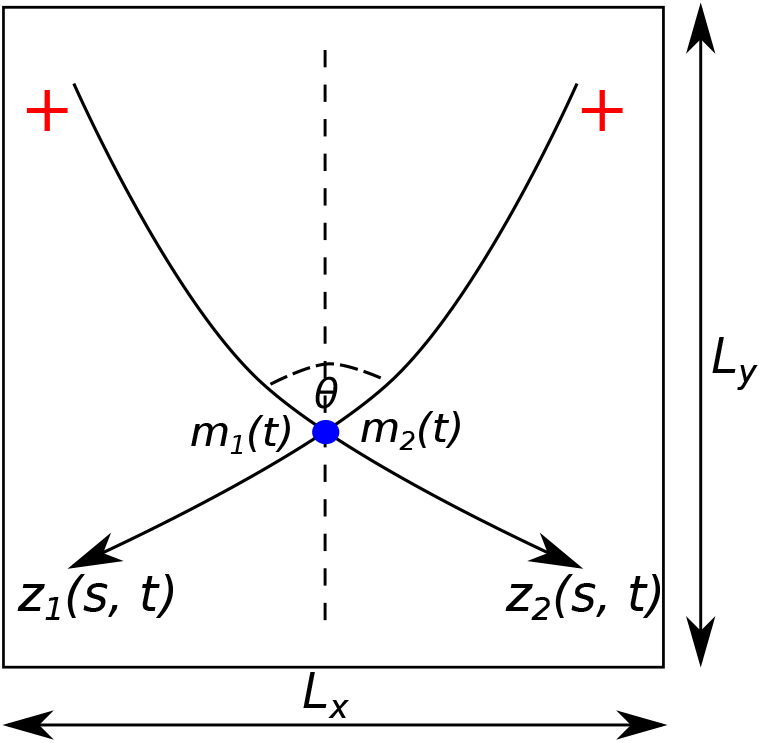
Schematic diagram of an actomyosin structure consisting of two filaments with an attached motor (blue dot). We characterise these structures by the mutual angle *θ* and motor positions *m*_1_ and *m*_2_.

Figure 3.4 illustrates two-filament simulations for both rigid and semi-flexible filaments. In the upper panel, the rigid filaments evolve symmetrically. As the motor traverses the filaments from the minus to plus ends, the filaments move and rotate such that their final position is a mirror image of the original. As reported by Lenz [9], this polarity-reversal symmetry causes the initial contraction and subsequent expansion to cancel. Principal stress arrows in the upper panel confirm this. The result is no net contraction for rigid filaments. However, the picture is different for semi-flexible filaments, as the lower panel reveals. When the motor begins to move, filament bending increases *θ*, increasing contraction in the *x*-direction. Subsequently, as the motor positions become favourable to expansion the angle between the filaments decreases (see the fourth image in the lower panel), decreasing the magnitude of expansion. Consequently, the semi-flexible filaments experience net contraction, providing evidence of the force asymmetry.

**Figure 3.4:**
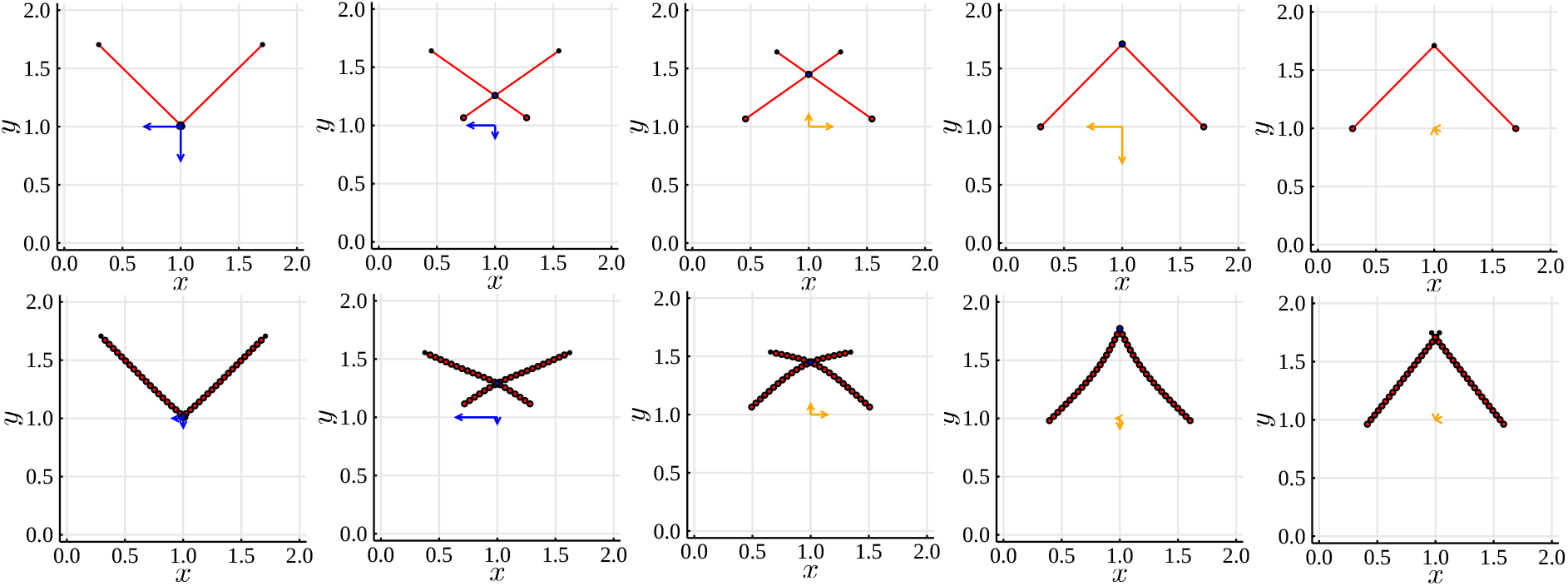
Two-filament simulations with initial motor positions *m*_1_ = *m*_2_ = 0, and *θ = π*/2. Upper panel: rigid actin filaments. Lower panel: Semi-flexible filaments. Results are presented (left–right) at *t* ∈ {0.04,1, 2.12, 3.12, 4}s. Arrows centred at (1,1) indicate the principal stress directions, and their lengths (given by the eigenvalues of ***σ*** represent the relative magnitude of stress. Blue arrows represent contraction, orange arrows represent expansion.

To verify this, we plot the bulk stress and *θ* versus time, for both rigid and semi-flexible filaments. The bulk stress results in Figure 3.5a confirm that rigid filaments experience no net contraction, because the magnitude of early contraction is equal to the magnitude of later contraction. The results in Figure 3.5b support this, where the angle *θ* for the initial contraction mirrors the angle for the subsequent expansion. In contrast, for semi-flexible filaments the structure experiences larger contractile than expansive stress. This is because filament bending generates an asymmetric pattern in *θ* with time, with a decrease as the motor approaches the plus ends. As a result, the semi-flexible filaments are unable to attain the large expansion that occurs towards the end of the rigid filament solution. This analysis confirms that a force asymmetry is a possible explanation for the bending-induced actomyosin contraction observed in §3.1.

**Figure 3.5:**
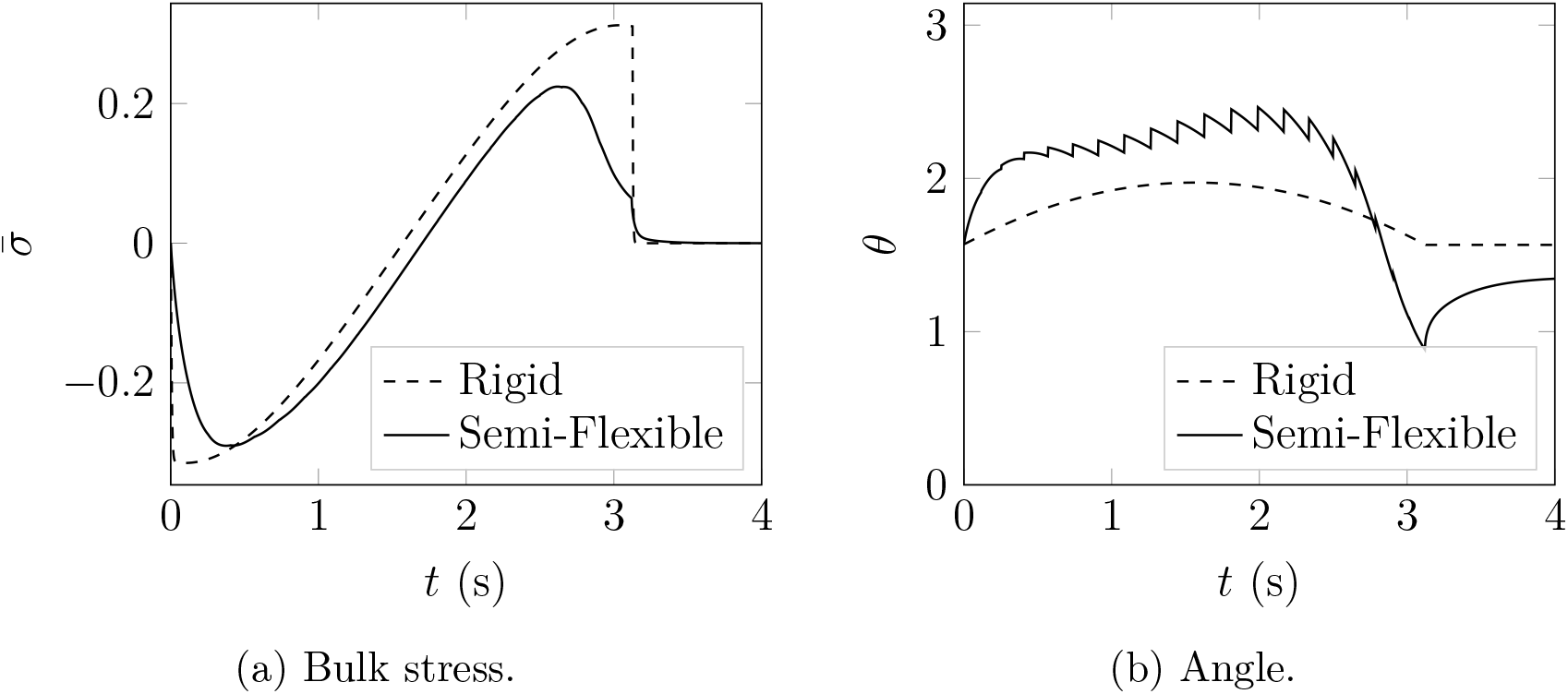
Bulk stress and *θ* versus time in two-filament simulations, for both rigid and semi-flexible filaments.

### 3.3 A Heuristic Index Predicts Stress Generated By Two-Filament-Motor Assemblies

Inspired by the previous results on the contraction of a two-filament-motor assembly, we propose a heuristic index that summarises the contractile potential of two filaments,

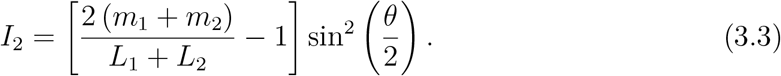

In (3.3), the left term in the brackets describes the length of the expansive and contractile branches, such that it is −1 if both motors are at the minus ends (contractile), and 1 if both motors are at the plus ends (expansive). To capture the effect of angle, the term in the right brackets is zero if *θ* = 0, and 1 if *θ* = *π*.

To confirm the effect of angle on contraction, we plot *I*_2_ (3.3) versus time in the two-filament simulations. In Figure 3.6, we multiplied *I*_2_ by a constant such that its minimum is equal to the minimum stress obtained in the simulation. The index accurately predicts the bulk stress in both simulations. Of particular note, *I*_2_ correctly predicts the loss of contraction with semi-flexible filaments, as Figure 3.6b shows. Combined with Figure 3.5b, this shows that filament bending generates contraction by influencing the angle between filaments, such that larger angles occur under contraction than under expansion.

**Figure 3.6:**
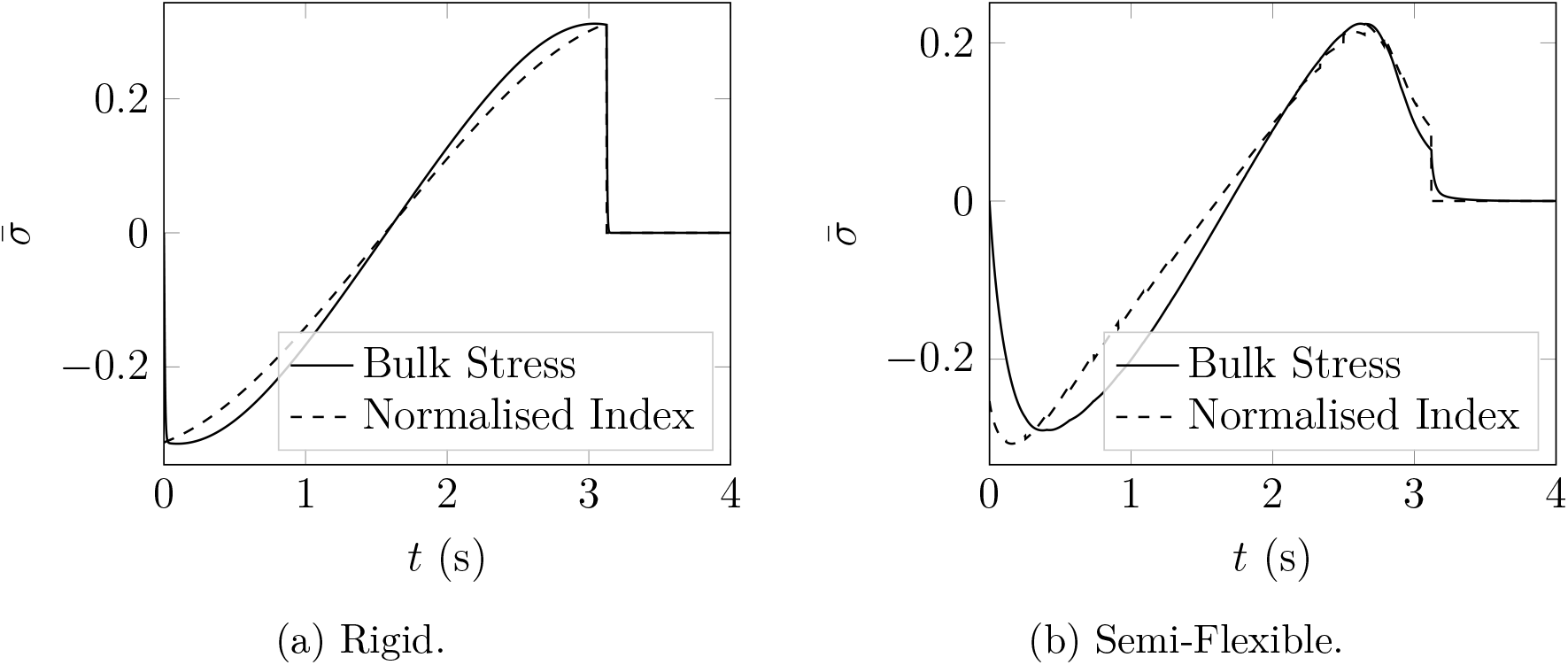
A comparison of the bulk stress and two-filament index (3.3) in the rigid and semi-flexible simulations. We normalised the two-filament index curve such that it has the same magnitude of contraction as the bulk stress.

To confirm the predictive ability of (3.3), we compute *I*_2_ for varying *m*_1_, *m*_2_, and *θ*. For each configuration, we compute one time step and compare the simulated bulk stress with (3.3). The results in Figures 3.7 and 3.8 show that the two-filament index *I*_2_ effectively captures the stress generated by two filaments. This is true if we hold *m*_1_ = *m*_2_ and vary *θ* (as in Figure 3.7), and if we hold *θ* constant and vary both *m*_1_ and *m*_2_ (as in Figure 3.8).

**Figure 3.7:**
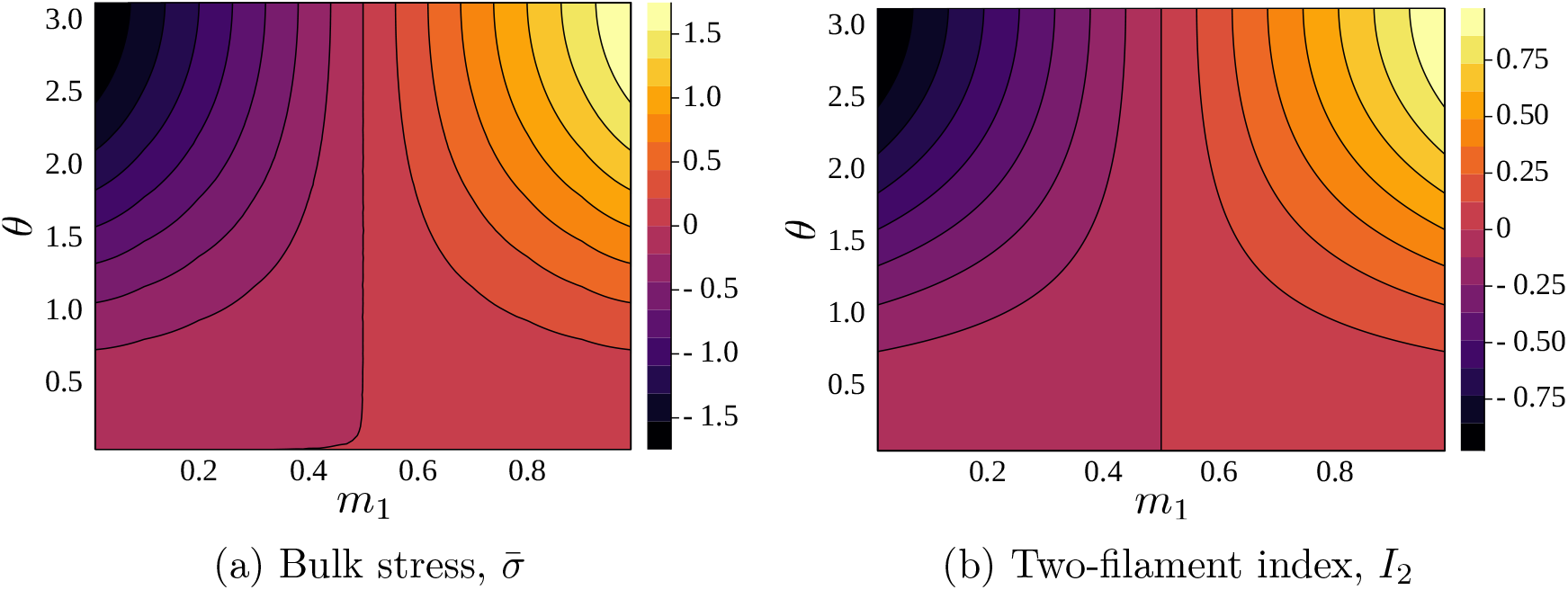
A comparison between bulk stress and the two-filament index (3.3), for one time step of a simulation with parameters as in Table 3.1, where Δ*t* = 2 × 10^−5^s. In these results, we set *m*_1_ = *m*_2_ and vary *θ* and *m*_1_.

**Figure 3.8:**
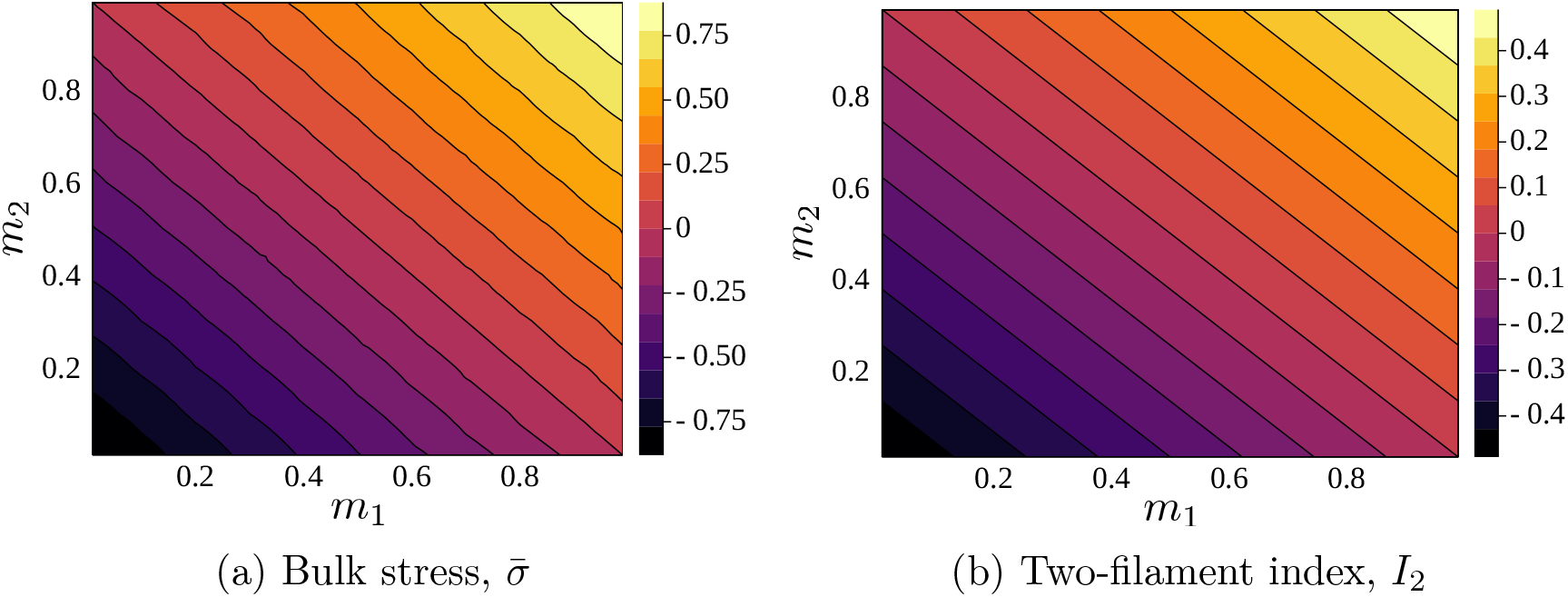
A comparison between bulk stress and the two-filament index (3.3), for one time step of a simulation with parameters as in Table 3.1, where Δ*t* = 2 × 10^−5^s. In these results, we set *θ* = *π*/2 and vary both *m*_1_ and *m*_2_.

Since the heuristic formula (3.3) is applicable to all two-filament assemblies, we will compare bulk stress with *I*_2_ in subsequent network-scale results.

### 3.4 Protein Friction Enables Network-Scale Contraction

Protein friction, either from cross-linking or filament contact, penalises relative motion where filaments overlap. Previous studies have suggested that intermediate cross-linker density maximises contraction [26, 30, 32, 42, 62, 63]. Without cross-linking, filaments move independently of each other, and are unable to generate collective contraction. However, strongly cross-linked networks generate large resistance to filament motion as myosin moves, which also inhibits contraction. To investigate this dependence using our model, we varied the protein friction drag coefficient, *λ*_pf_, and computed ten simulations with each parameter value. Results from these simulations are shown in Figure 3.9.

**Figure 3.9:**
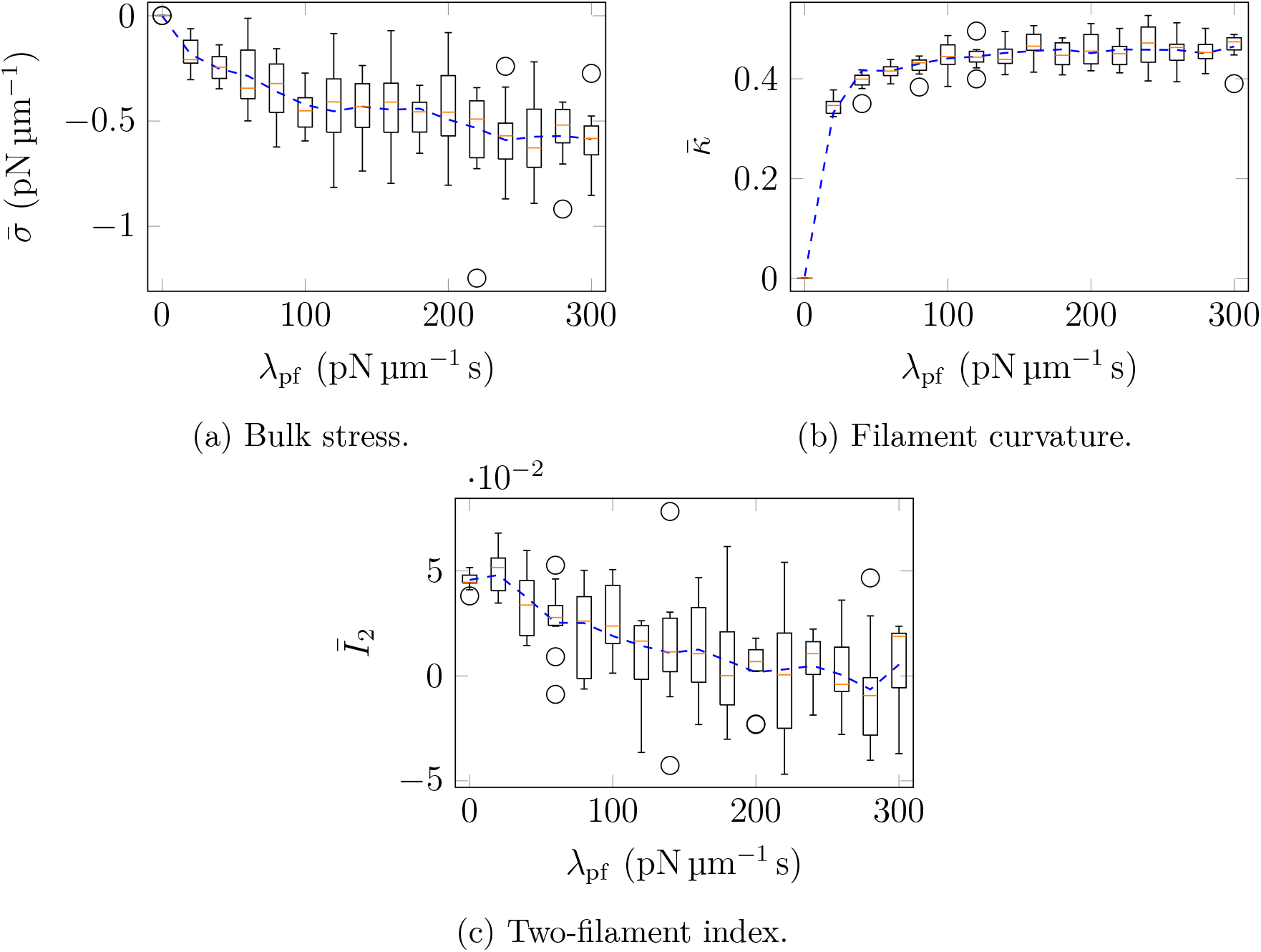
Simulation results for the effect of the protein friction coefficient, *λ*_pf_, on time-averaged quantities of interest. Box plots represent ten random simulations for a given parameter, orange bars indicate median values, and dashed curve is the mean data, smoothed with a Savitsky–Golay filter.

Figure 3.9a shows the relationship between *λ*_pf_ and bulk stress. As expected, networks become more contractile as *λ*_pf_ increases from zero. Although the precise value of the protein friction coefficient for actin filaments is unknown, Ward et al. [49] suggests protein friction due to filament contact of approximately *λ*_pf_ = 30 pN μm^−1^ s. Estimating *λ*_pf_ based on the cross-linker *α*-actinin yields approximately *λ*_pf_ = 20 pN μm^−1^ s (see Supplementary Material). Both values are sufficient to demonstrate contractile bias. Subsequent increases in *λ*_pf_ beyond these values incur diminishing returns, such that contractility becomes stable after approximately *λ*_pf_ = 200 pN μm^−1^ s. We do not observe a U-shaped curve in stress with *λ*_pf_, possibly because the relative sparseness of our simulated networks does not enable sufficient connectivity to restrict contraction.

Plots of the time-averaged curvature and *I*_2_ in Figures 3.9b and 3.9c respectively demonstrate that contraction correlates with increased curvature and decreased *I*_2_. An important finding is that filament bending does not occur in the absence of protein friction. This is because protein friction supplies resistance to motion at specific points along the filament. Without this drag, the filament will tend to adopt the energetically preferable straight configuration. Therefore, protein friction is essential to generating contraction. Furthermore, only a small increase in filament bending is attainable by increasing the protein friction coefficient beyond the biologically-feasible value of *λ*_pf_ = 20 pN μm^−1^ s.

### 3.5 Viscous Friction Inhibits Contraction

The viscous drag coefficient *λ_a_* represents drag between actin filaments and structures external to the network. This can arise from drag between the filaments and the cytoplasm, or drag between filaments and a dense, homogeneous background network that interacts uniformly with the simulated filaments. Increasing *λ_a_* thus corresponds to increasing cytoplasm viscosity, or increasing the network density. *In vitro* experiments by Murrell and Gardel [16] showed that increasing adhesion between actomyosin networks and the membrane inhibits contraction. We suggest that increased membrane adhesion corresponds to an increase in drag coefficient in our model, because both restrict filament motion. For these reasons, we are interested in how contractility depends on *λ_a_*.

We varied *λ_a_* and performed ten simulations for each parameter value. These results are shown in Figure 3.10. As predicted by experiments, network contractility increases as we decrease *λ_a_*. Interestingly, Figures 3.10b and 3.10c show that this increased contraction does not correspond to an increase in filament curvature or decrease in the two-filament index. Instead, a possible explanation is that increasing *λ_a_* increases resistance to actin filament movement. When myosin motors exert forces on the network, a larger proportion is used to overcome drag as *λ_a_* increases. This inhibits the ability of myosin motors to remodel the network, and this slower remodelling results in decreased contraction.

**Figure 3.10:**
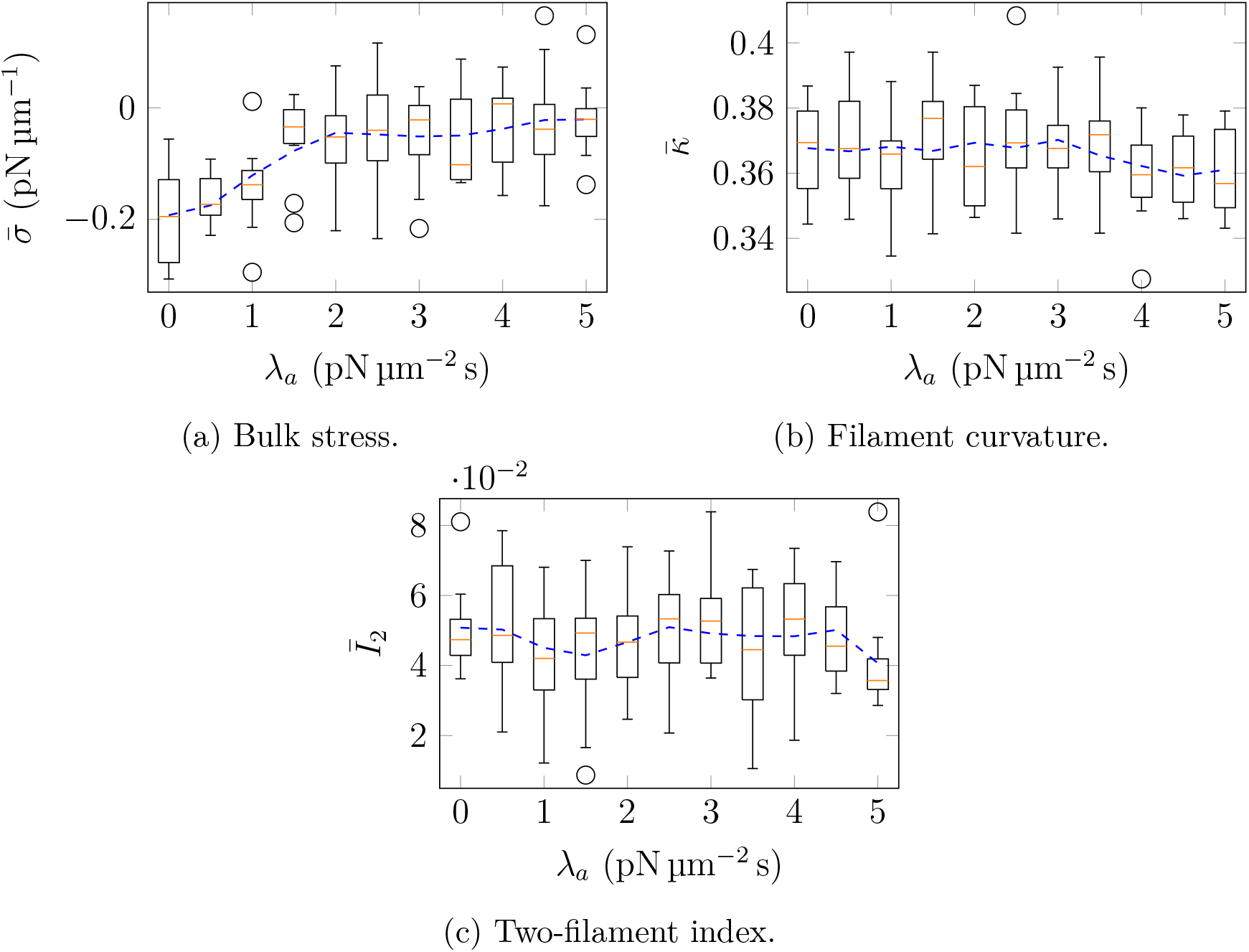
Simulation results for the effect of actin–background drag coefficient, *λ_a_*, on time-averaged quantities of interest. Box plots represent ten random simulations for a given parameter, orange bars indicate median values, and dashed curve is the mean data, smoothed with a Savitsky–Golay filter.

### 3.6 Myosin Unbinding Has Negligible Effect on Contractility

Myosin motor unbinding is another feature of our model that might influence contractility. In our simulations, motor unbinding is governed by Bell’s law. All motors that have not reached the end of a filament unbind with a rate that depends on the spring force on the motor, and the reference off-rate, *k*_off,*m*_. To investigate how this off-rate affects contractility, we computed a series of simulations with varying *k*_off,*m*_, and present results in Figure 3.11.

**Figure 3.11:**
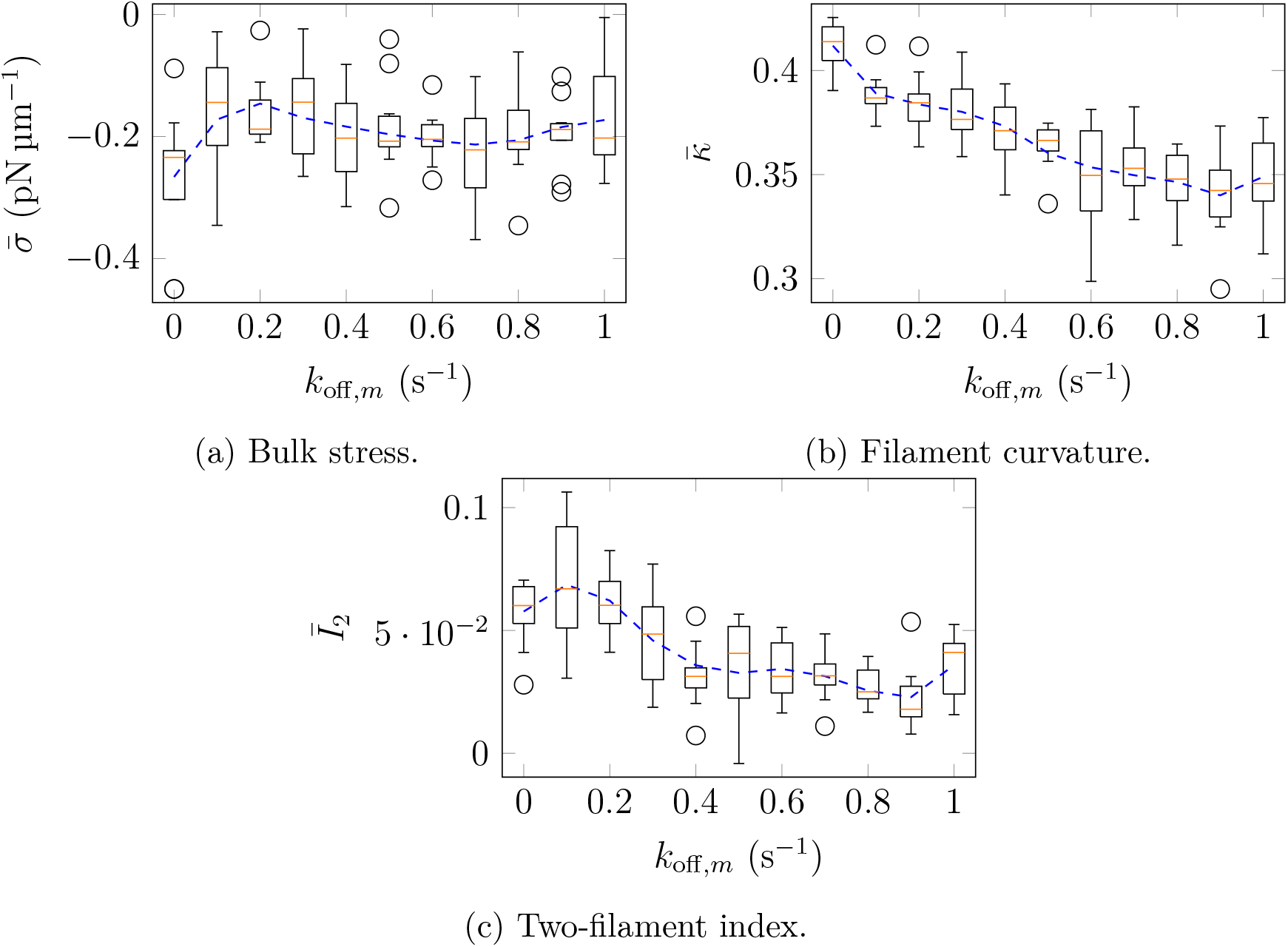
Simulation results for the effect of myosin motor reference off-rate, *k*_off,*m*_, on time-averaged quantities of interest. Box plots represent ten random simulations for a given parameter, orange bars indicate median values, and dashed curve is the mean data, smoothed with a Savitsky–Golay filter.

Overall, the reference off-rate has no discernible effect on stress. However, Figures 3.11b and 3.11c suggest that the means of generating contraction changes as *k*_off,*m*_ changes. A possible explanation is that *k*_off,*m*_ governs the expected time for which a motor remains attached to the filaments. For example, lower values of *k*_off,*m*_ enable motors to remain attached to actin filaments for longer time. Highly-persistent motors have longer time to initiate bending, and therefore curvature increases as *k*_off,*m*_ decreases (see Figure 3.11b).

However, such persistent motors also walk further towards the plus ends, increasing *I*_2_ (see Figure 3.11c). As shown in §3.3, motor positioning closer to the plus ends is favourable for expansion. The competing effects of filament bending and motor position enable disordered networks to generate similar contractile stress for all reference motor off-rates tested.

We also tested whether the force-dependence introduced by Bell’s law influences contractility. To do this, we performed ten simulations with both rigid and semi-flexible filaments, and compare the time-averaged bulk stress results with the default simulations in Figure 3.1. Simulation results with force-independent unbinding are given in Figure 3.12. Since the results are similar to those in Figure 3.1, we conclude that the force-dependence introduced by Bell’s law has insignificant effect on the network dynamics, for the biophysically-realistic parameters investigated.

**Figure 3.12:**
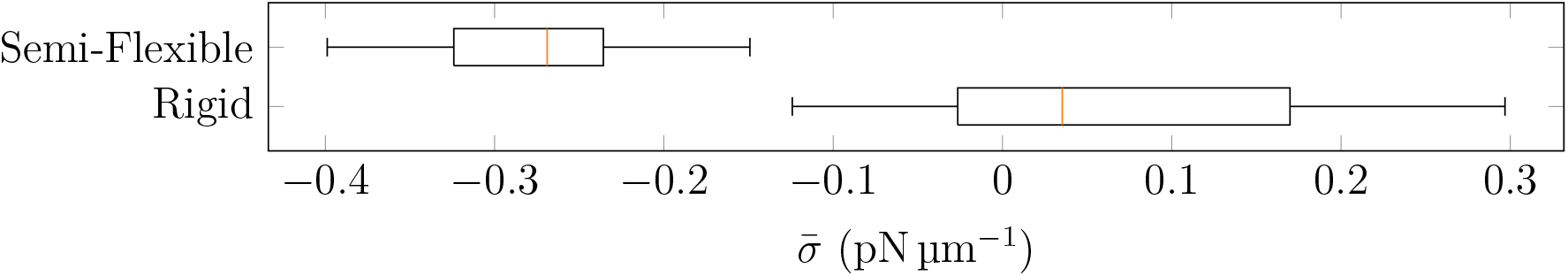
Box plots of time-averaged bulk stress 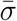 in ten semi-flexible simulations and ten rigid simulations, using default parameters with force-independent myosin unbinding (no Bell’s law). Box plots represent ten random simulations for a given parameter, and orange bars indicate median values.

### 3.7 Actin Filament Turnover Enables Persistent Contraction

In biological cells, actin filament turnover is an important process that enables sustained contraction. Turnover refers to the exchange of proteins with the background cytoplasm, and introduces randomness. Without turnover, actomyosin networks have been shown to lose contractility over time [7, 12, 17, 28, 42, 46]. To investigate whether our model replicates this behaviour, we varied the actin filament turnover rate, *k*_off,*a*_, and present results for the simulated stress in Figure 3.13. Time-averaged stress results show increased contraction as we increase actin turnover rate. In support of this, Figures 3.13b and 3.13c show that increased actin turnover corresponds to a decrease in mean integrated filament curvature, and the two-filament index shows bias towards expansive configurations.

**Figure 3.13:**
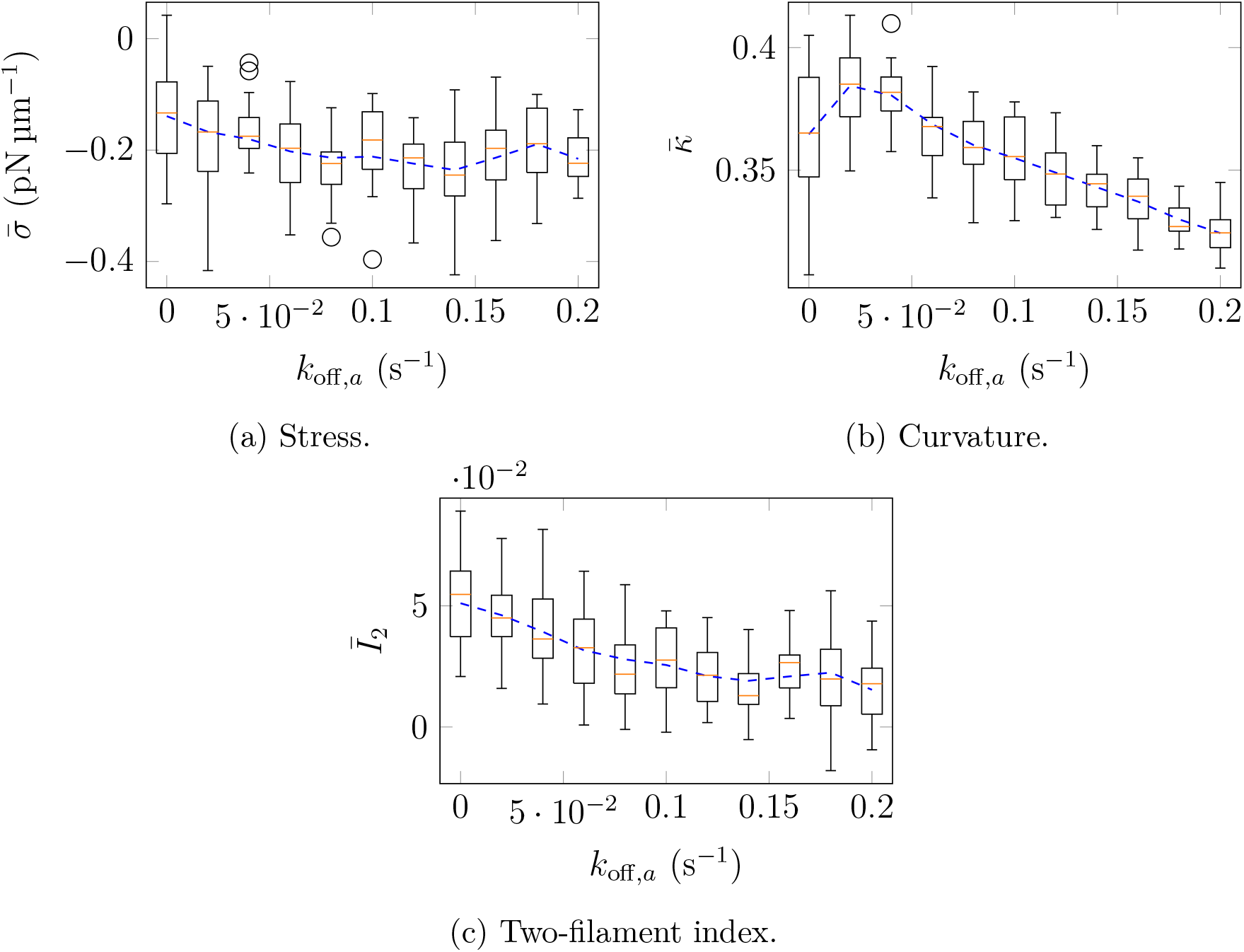
Simulation results for the effect of actin turnover rate, *k*_off,*a*_, on time-averaged quantities of interest. Box plots represent ten random simulations for a given parameter, orange bars indicate median values, and dashed curve is the mean data, smoothed with a Savitsky–Golay filter.

To investigate the time-dependence of contractile stress with and without turnover, we plot the mean bulk stress in the ten simulations versus time for *k*_off,*a*_ = 0 s^−1^ (no turnover) and *k*_off,*a*_ = 0.2s^−1^ (fast turnover). With no turnover, there is a loss of contractility as time progresses (see Figure 3.14a), whereas no trend occurs with fast turnover. Since both networks in Figure 3.14 show similar contractile stress at *t* = 0, the results in Figure 3.13a occur because the network loses contractility if there is no turnover, decreasing time-averaged stress 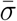.

**Figure 3.14:**
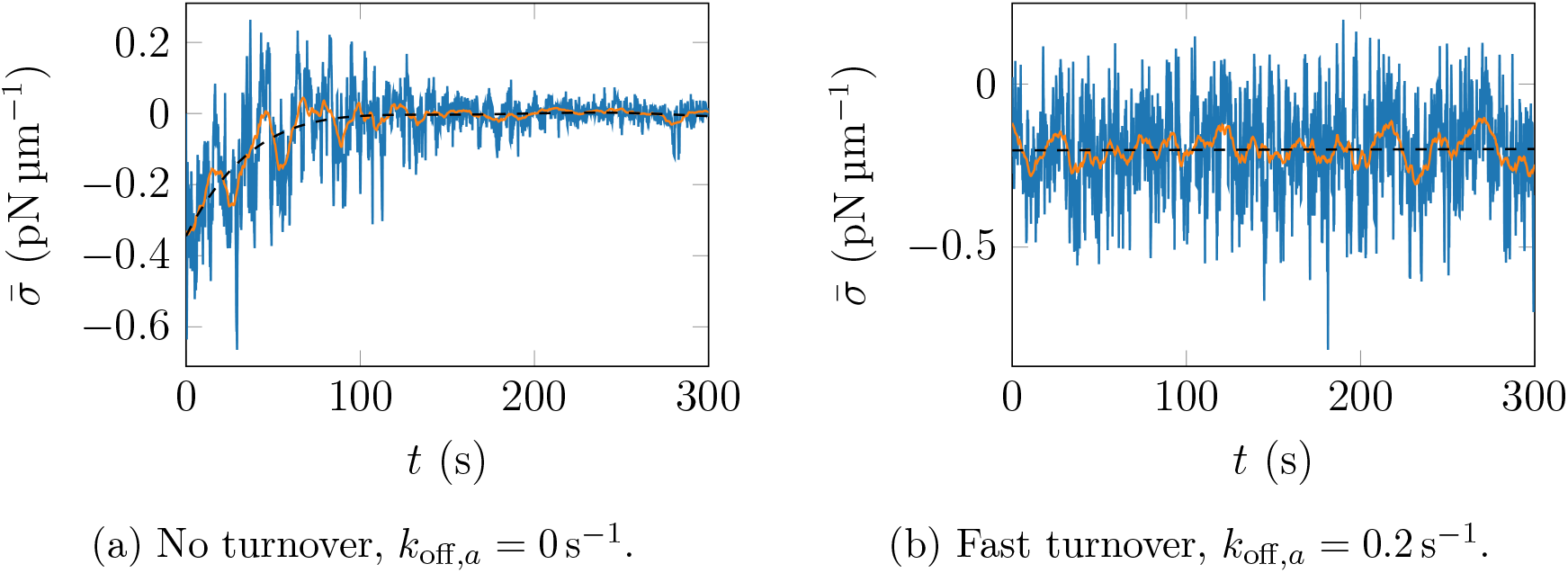
Mean bulk stress across ten simulations versus time. The blue curve indicates mean data across ten simulations. The orange curve is a moving average of the mean data, with a window width of 10 s, and the black curve is a fit to the mean data. All parameters as given in Table 3.1, except for actin turnover rate.

Previous studies have shown that loss of contraction in the absence of turnover is associated with pattern formation in the network. This involves filaments aggregating in asters [64] or bundles [65], after which they do not move under molecular motor activity. To investigate whether pattern formation occurs in our simulations, we computed the distance between all pairs of nodes on different filaments. If the distribution of these distances differs from the expected distribution for two random points in a square, we conclude that filaments have aggregated. An example comparison of these distance distributions at 300 s, and the corresponding network images, are provided in Figure 3.15. With no turnover, there are two peaks in the distribution of distances that are not predicted by the theoretical distribution. In contrast, the distance distribution closely matches the theoretical distribution in the simulation with fast turnover, *k*_off,*a*_ = 0.2 s^−1^. This provides evidence that actin filaments aggregate with no turnover, indicating pattern formation as expected. Fast turnover prevents this pattern formation by introducing randomness to filament positions, enabling persistent contraction. Similar distributions occur across all simulations, a complete summary of which is given in the Supplementary Material.

**Figure 3.15:**
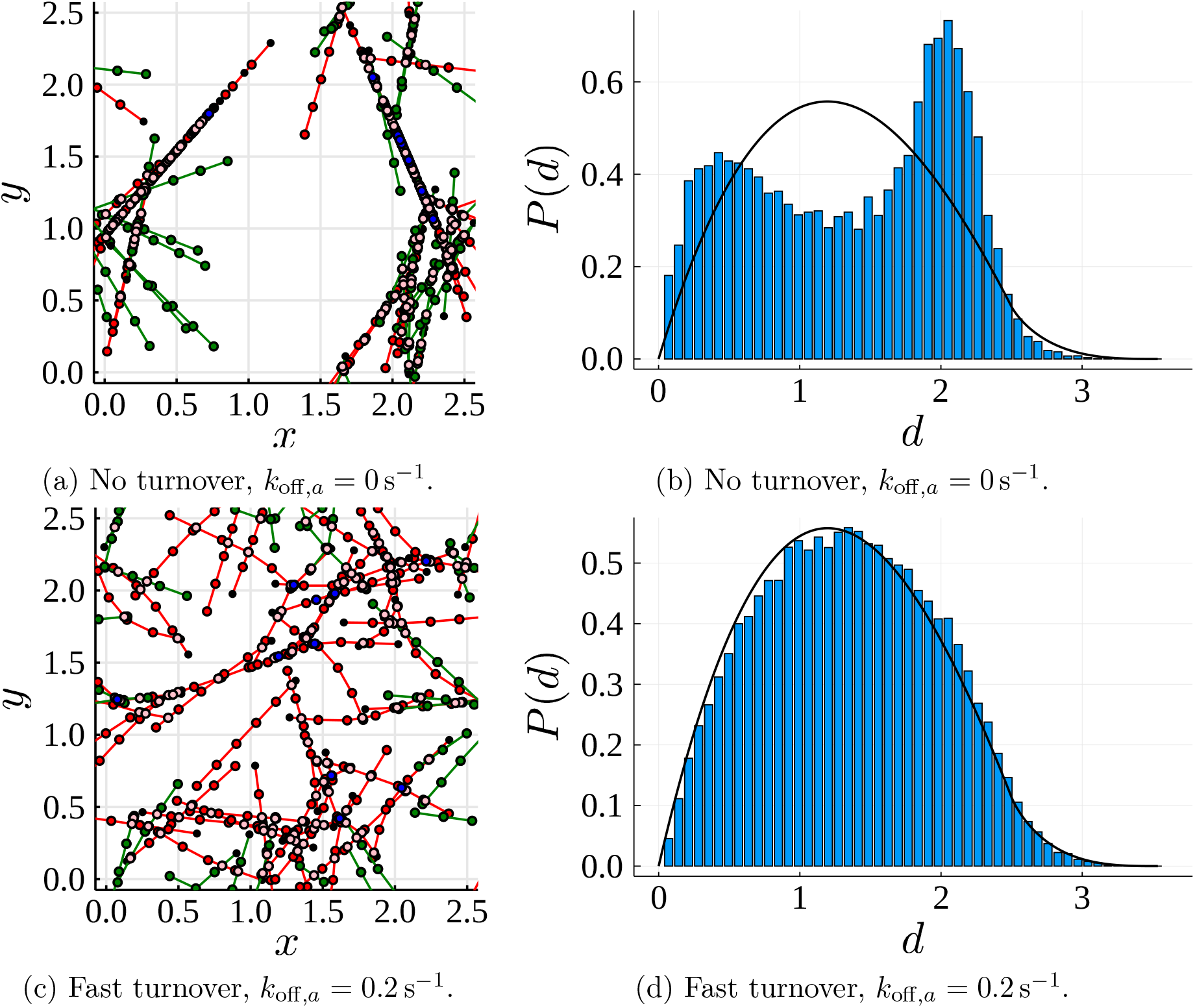
Network configurations and probability distributions of the distances between plus ends of all pairs of actin filaments in the network at *t* = 300 s. Blue bars indicate simulation results, and the black curve is the theoretical distribution for the distance between two random points in a square [66].

### 3.8 Actomyosin Network Contraction Without Molecular Regulation Does Not Exhibit Periodic Pulsation

Interestingly, periodic or pulsed contraction has been observed in experiments and simulations with filament turnover [17, 42, 67, 68]. Some authors have suggested that biochemical signals external to the network are responsible for this pulsation [67, 68]. However, recent work by Yu et al. [69] showed that pulsation might be an inherent result of actomyosin mechanics, caused by actin treadmilling or severing. As Figure 3.14a shows, stress rises and falls in our simulations with or without turnover, indicating pulse-like behaviour. To investigate whether solutions with turnover have a characteristic period of pulsation, we computed a simulation with default parameters for 30 minutes of simulated time. Plotting the autocorrelation of the stress signal then enables us to determine whether a characteristic period exists. These results are shown in Figure 3.16. Autocorrelation compares original stress signal and a time-delayed version, and returns the correlation coefficient at a function of the time delay. If stress generation is periodic with period *T*, we would see peaks in the autocorrelation at all multiples of *T*. In Figure 3.16, no such peaks appear in the first ten minutes of the solution. Therefore, although our results show oscillations in contractile stress, these oscillations are aperiodic. Consequently, in our simulations contractile pulses arise due to random variations in the network, without a systematic mechanical origin.

**Figure 3.16:**
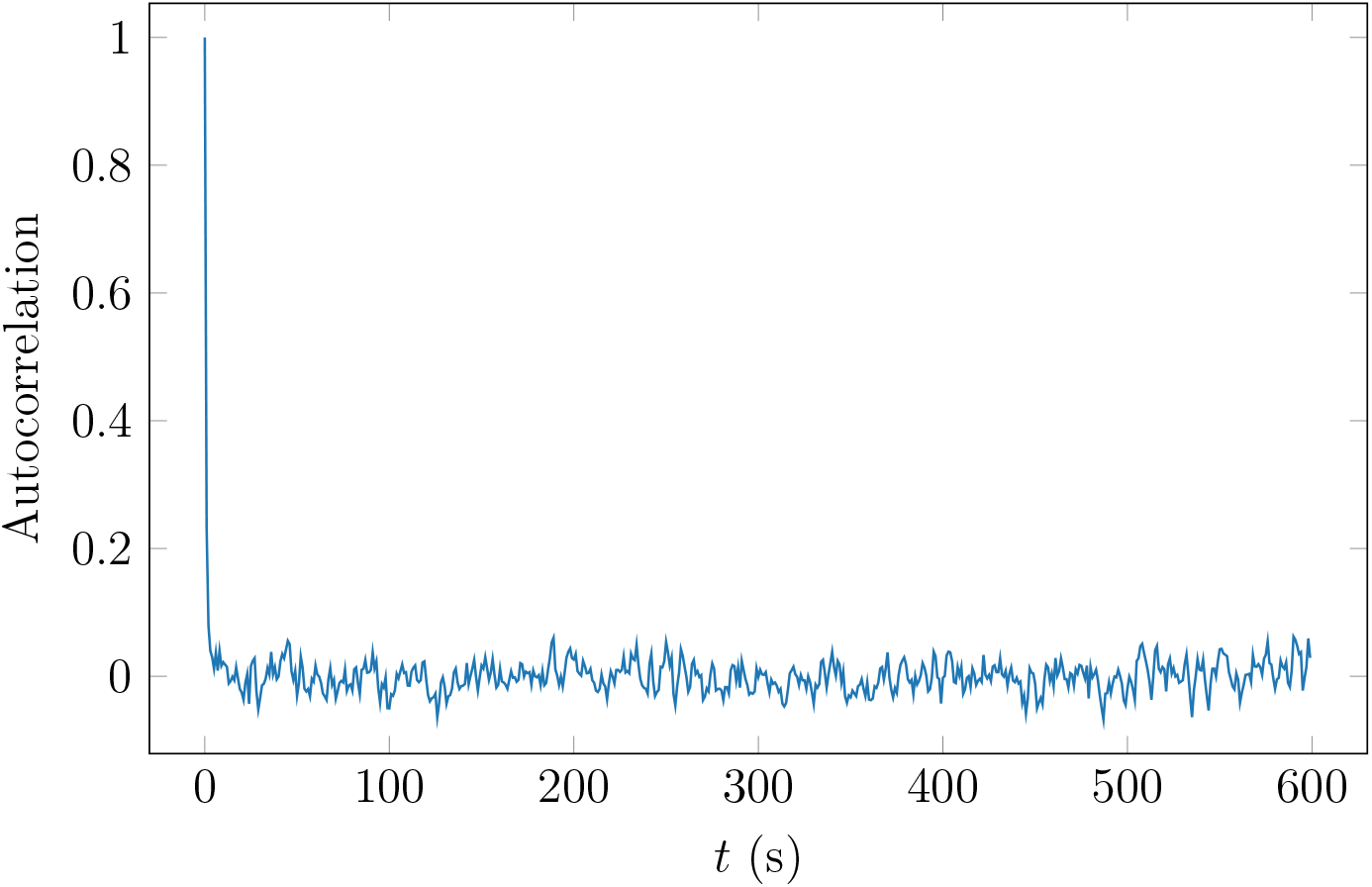
Autocorrelation function for the stress signal in a simulation with default parameters from Table 3.1, except for *T* = 1800 s.

Our findings extend the results of Belmonte, Leptin, and Nédélec [17], who used visual inspection of simulations to show that pulsation occurs in networks with turnover. Our results are consistent with observations that pulsation occurs due to biochemically-regulated, periodic formation of actomyosin networks [67, 68], and not necessarily periodic stress generation within the networks. Observing periodic mechanical behaviour would require additional assumptions to those in our model, for example actin treadmilling or severing, which Yu et al. [69] showed to be necessary for pulsed contraction in the absence of biochemical regulation.

## 4 Conclusion

Contraction of disordered actomyosin networks is essential to biological cell function. Since the origins of this contraction are not yet fully understood, scientists have worked to build an inventory of possible contraction mechanisms. In this study, we investigated the hypothesis that protein friction, arising from cross-linking or solid friction between actin filaments, enables contraction of networks consisting of semi-flexible actin filaments. We achieved this by developing an agent-based mathematical model for two-dimensional actomyosin networks. Using a variational, gradient-flow like formulation of the system of force-balance equations, our model provides a new method of quantifying network stress. Numerical simulations confirmed that actin filament bending generates a force asymmetry that biases contraction over expansion in random networks. Importantly, network-scale bending is only possible with protein friction, making protein friction crucial to generating contraction.

To understand the bending-induced force asymmetry at the microscopic scale, we simulated the simplest actomyosin structure consisting of a single myosin motor bound to two actin filaments. For both rigid and semi-flexible filaments, the contractile force depends on the motor relative positions, and the angle between the two filaments. As the motor moves from the minus to the plus ends, semi-flexible filaments generate a wider angle than rigid filaments. Since these wider angles are more conducive to contraction, our microscopic simulations showed that filament bending induces contractile bias at the microscopic scale. Furthermore, this confirmed that bending forces are sufficient to generate contraction, without the need to invoke filament buckling.

Our simulations also confirmed previous experimental and theoretical results that filament turnover is required to sustain contraction. Although actin bending and protein friction generate contraction, without turnover the filaments aggregate and form patterns, after which the network loses contractility. In our simulations, introducing turnover generates a more random spatial distribution of filaments, and enables the network to sustain contractility. However, in many cell types actin filaments can form contractile actomyosin bundles such as stress fibres, which are aggregated structures that sustain and mediate contractility [70]. An important extension to our work will be to identify the minimal mechanisms that enable self-organisation and persistence of such bundles, even in networks with fast turnover. We plan to extend the modelling framework developed here to tackle this problem in future work.

## Author Contributions

A. K. Y. T. and D. B. O. designed the research and developed the mathematical model. A. K. Y. T. performed the numerical simulations, analysed the data, and wrote the manuscript, with guidance from D. B. O and A. M.

## Acknowledgements

The authors acknowledge funding from the Australian Research Council (ARC) Discovery Program (grant number DP180102956), awarded to D. B. O. and A. M.

## A Mathematical Model Derivation

We develop and implement an agent-based mathematical model for two-dimensional actomyosin networks. We represent actin filaments as finite-length curves in 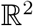, and to track their position introduce the variables 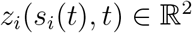 for *i* = 1,...,*N_a_*, where *N_a_* is the number of semi-flexible actin filaments. These represent the physical position of the actin filament, parameterised by the arc length *s_i_*(*t*) ∈ [0, *L_i_*], where *L* is the length of the *i*-th actin filament. We consider a simplified representation of myosin motors as dumbbells that behave like stiff linear springs. The two ends of the dumbbell represent motor ‘heads’ that bind to actin filaments and exert forces. To track motor head positions, we define the variables *m_ik_*(*t*) ∈ [0, *L_i_*], for *k* = 1,..., *N*_m_, where *N_m_* is the number of myosin motors. These are the positions (measured from the minus end) of the *k*-th myosin motor along the actin filament with index *i*, to which it is bound. The derivation of our model in a time-discrete context then involves constructing an energy functional that depends on the degrees of freedom *z_i_* and *m_ik_*. At each time step, the solution is given by the minimiser of this functional, and advancing in time enables us to simulate network evolution. We solve the model on a two-dimensional domain with periodic boundary conditions, such that the network evolves on the surface of a torus.

### A.1 Energy Functional

We write the mathematical model in a time-discrete context in terms of an energy functional that depends on the degrees of freedom *z_i_*(*s_i_*(*t*),*t*), and *m_ik_*(*t*). This functional consists of the potential energy contribution of each mechanical feature in the model, and at each time step the network evolves to minimise this energy. In abstract terms, the total network energy consists of

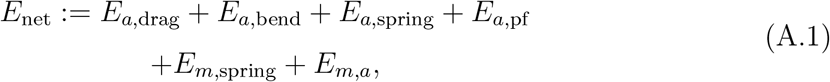

where the subscripts *a* and *m* refer to actin and myosin respectively. Below, we outline the meaning and mathematical description of each term in (A.1).

We assume that viscous drag with a background medium resists motion of the actin filaments. We then obtain the actin drag energy contribution

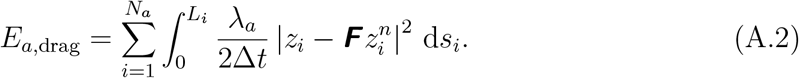

In (A.2), *λ_a_* is the coefficient of viscous drag for actin–background interactions, and is similar to the damping term *λ* in the Langevin equation. The vector 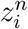 represents filament positions at the previous time step, where Δ*t* is the time step size. To account for stretching and rotation of the domain, we multiply 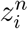 by the deformation gradient tensor

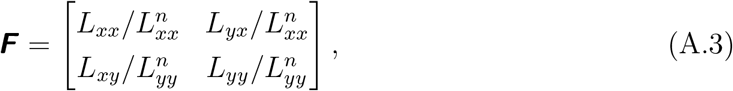

which ensures both *z_i_* and 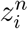 are represented in the current spatial co-ordinates. In network-scale simulations, this drag term represents hydrodynamic drag with the background cytoplasm. An alternative interpretation of viscous drag is to assume that the simulated network is a subset of a dense, homogeneous, cross-linked network of filaments.

Since filaments are semi-flexible, we also include the contribution of elastic potential energy due to bending. This is given by

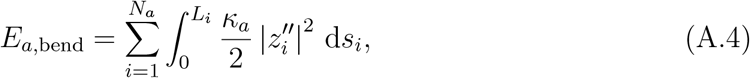

where *κ_a_* is the flexural rigidity, assumed constant for all actin filaments. The third term in (A.1), *E*_*a*,spring_, is the energy associated with local longitudinal extension of actin filaments. According to Hooke’s law, after summing the contributions of all filaments, it is given by

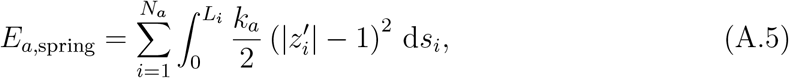

where we *k_a_* specifies the actin filament stiffness, and is assumed to be the same for all filaments. Note that in the context of our model we regard (A.5) as a penalising potential with large coefficient *k_a_* in order to model actin filament in-extensibility and to be able to regard *s_i_* as arc-length parametrisation.

Protein friction between actin filaments also contributes to the energy functional. In our model, we represent this as a viscous drag contribution that acts point-wise at intersections between actin filaments. This viscous force can arise due to contact friction between overlapping filaments [49], or as the macroscopic effect of abundant cross-linkers that undergo turnover [38]. The energy contribution due to protein friction is

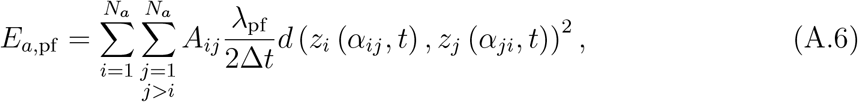

where *λ*_pf_ is the protein friction drag coefficient. In (A.6), *A_ij_* is a binary variable such that *A_ij_* = 1 if filaments *i* and *j* intersect and no motor is bound to both filaments, and *A_ij_* = 0 otherwise. We also define *d*(*z*_1_, *z*_2_) to be the shortest physical distance between two points 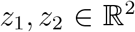 or their periodic translations, enabling us to account for periodic boundary conditions. Finally, *α_ij_* ∈ [0,*L_i_*] is the position along filament *i* at which the intersection with filament *j* occurs, and ensures that protein friction drag is applied point-wise at these intersections.

The final two terms in (A.1) model the effects of myosin motors. In the same way as we account for F-actin inextensibility, we use the penalising potential

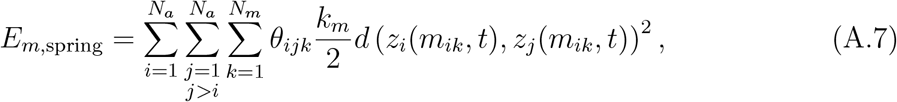

to model myosin inextensibility. Here *k_m_* is the myosin motor spring constant which we take as very large, and *θ_jk_* is a binary variable such that *θ_jk_* = 1 if myosin motor *k* is attached to filaments *i* and *j*, and *θ_jk_* = 0 otherwise. The final term in (A.1) describes interactions between filaments and motors. We assume that myosin obeys a linear force–velocity relation, such that positions evolve according to

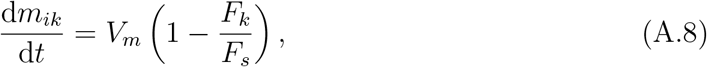

where *V_m_* is the load-free myosin motor velocity, *F_s_* is the motor stall force, and *F_k_* is the force on the *k*-th myosin motor. To reproduce this term (A.8) as the variation of an energy for discrete time, we introduce a linear term for the load-free velocity, and a quadratic term with the same scaling as the drag terms above for the linear velocity reduction due to motor loading. The energy then reads

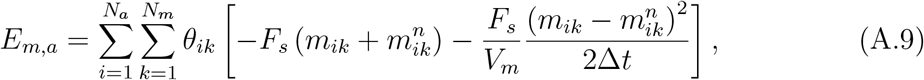

where *θ_ik_* is a binary variable such that *θ_ik_* = 1 if motor *k* is attached to filament *i*, and *θ_ik_* = 0 otherwise. This completes the description of all terms in the network energy functional. In lieu of a detailed description of the numerical scheme used to solve the model, we make available on Github the Julia code used to perform the simulations.

### A.2 Stochastic Filament and Motor Turnover

We simulate random actin filament turnover and myosin motor unbinding. Given an off-rate *k*_off_, the probability of turnover or detachment in a given time step according to an exponential distribution is

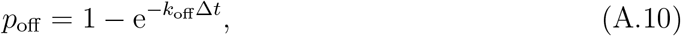

where Δ*t* is the time step size. We assume that the turnover rate for actin filaments, *k*_off,*a*_, is constant and the same for each filament. At each time step, we use a pseudo-random number generator to simulate whether each filament will turn over. To maintain constant filament density, we immediately replace filaments that turn over with new ones at random positions and orientations. If a filament turns over, we also assume that any myosin motor attached to the filament automatically unbinds.

In contrast, we assume that the unbinding rate for myosin motors depends on the force it experiences. According to Bell’s law, the force-dependent unbinding rate is given by

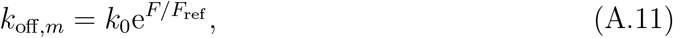

where *k_0_* is the reference off-rate for unloaded motors, and *F*_ref_ is a reference force. The force to which the *k*-th motor is subject is the variation of the penalising potential (A.7) and given by a Hooke’s law, where motors are assumed to be linear springs with equilibrium length zero. This yields *F_k_* = *k_m_d*(*z_i_*(*m_ik_, t*) − *Z_j_*(*m_jk_,t*)), where *i* and *j* are the indices of the two filaments to which the motor attaches, such that the distance term measures the motor length. Like the actin filaments, we maintain constant myosin motor density throughout the simulation by assuming that an unbound motor is immediately replaced with a new one at a random filament intersection.

### A.3 Parameters

We performed network simulations in the main text with a set of default parameters. These parameters are listed in Table A.1. Additional information on the derivation of some parameters is provided below.

**Table A.1:** Default parameters for actomyosin network simulations.

| Parameter | Symbol | Value | Units | Source |
| --- | --- | --- | --- | --- |
| Longitudinal stiffness (actin) | $k_a$ | 1000 | pN | [28] |
| Longitudinal stiffness (myosin) | $k_m$ | 1000 | pN $\mu\text{m}^{-1}$ | [28] |
| Actin filament flexural rigidity | $\kappa_a$ | 0.073 | pN $\mu\text{m}^2$ | [52] |
| Equilibrium actin filament length | $L_a$ | 1 | $\mu\text{m}$ | [10, 18, 40] |
| Actin–cytoplasm drag coefficient | $\lambda_a$ | 0.05 | pN $\mu\text{m}^{-2} \text{s}$ | [45, 53, 54] |
| Protein friction drag coefficient | $\lambda_{\text{pf}}$ | 30 | pN $\mu\text{m}^{-1} \text{s}$ | [49] |
| Myosin stall force | $F_s$ | 5 | pN | [12, 55, 56] |
| Myosin free-moving velocity | $V_m$ | 0.5 | $\mu\text{m} \text{s}^{-1}$ | [12, 56, 57] |
| Actin filament turnover rate | $k_{\text{off}}$ | 0.04 | $\text{s}^{-1}$ | [21, 43, 58] |
| Myosin reference off-rate | $k_0$ | 0.35 | $\text{s}^{-1}$ | [59, 60] |
| Myosin reference unbinding force | $F_{\text{ref}}$ | 12.6 | pN | [61] |
| Number of actin filaments | $N_a$ | 50 | [–] | Assumption |
| Number of myosin motors | $N_m$ | 10 | [–] | Assumption |
| Domain width | $L_{xx}, L_{yy}$ | 2.5 | $\mu\text{m}$ | Assumption |
| Simulation duration | $T$ | 60 | s | Assumption |

#### Longitudinal Stiffnesses *k_a_, k_m_*

We assume that actin filament segments and myosin motors are stiff entities, and following Stachowiak et al. [28] use *k_a_* = *m_m_* = 1000 pN μm^−1^. By inspection, these values are sufficiently large to ensure filament segments and myosin motors experience negligible extension.

#### Actin Filament Length

Actin filament length depends on cell type and function, and can vary across experiments. Since our modelling follows Hiraiwa and Salbreux [10] and Dasanayake, Michalski, and Carlsson [40], we adapt estimates from these authors. Dasanayake, Michalski, and Carlsson [40] use *L_a_* = 2 μm, whereas Hiraiwa and Salbreux [10] use *L_a_* = 0.1–1 μm. Experimental measurements of fission yeast by Kamasaki, Osumi, and Mabuchi [18] give *L_a_* = 0.6 μm, and Stachowiak et al. [28] use *L_a_* = 1.3 μm. Based on this data, a reasonable estimate for our model is *L_a_* = 1 μm.

#### Actin–background drag coefficient, *λ_a_*

Since the actin–background drag coefficient is difficult to estimate, we assume *λ_a_* = 0.05 pN μm^−2^ s in network simulations. This value is small enough that actin–background drag has minimal effect on the network. For an experimental justification of this parameter, we modify the formula from Berg [53] that describes the frictional drag coefficient of an ellipsoidal object moving at random through an incompressible viscous medium. The formula is

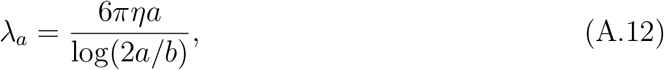

where *η* is the viscosity of the medium (in this case the cytoplasm), *a* is the semi-major axis length (*i.e*. half the filament length), and *b* is the semi-minor axis length (*i.e*. the actin filament radius). Based on Oelz et al. [54], we modify (A.12) and use the formula

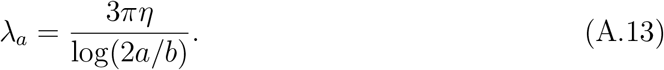

We assume filaments have the constant length *L_a_* = 1 μm, and thus *a* = 0.5 μm. The actin filament has a diameter of 7nm [71, 72], such that the radius is *b* = 0.0035μm. The drag coefficient *λ_a_* = 0.05pN s μm^−2^ then corresponds to *η* = 0.03pN s μm^−2^, which is approximately 30 times the viscosity of water. According to a recent review, the cytoplasm viscosity is 2–50 times that of water [73], and therefore our estimate is reasonable.

#### Protein Friction Drag Coefficient, *λ*_pf_

We estimate the protein friction drag coefficient using experimental work by Ward et al. [49] on sliding friction between F-actin filaments. Given a pulling velocity of 0.2 μms^−1^, they obtain a frictional force of approximately 6 pN, suggesting that *λ*_pf_ = 30 pN μm^−1^ s.

Under the alternative interpretation of protein friction as the macroscopic effect of abundant, transient cross-linkers, we can estimate *λ*_pf_ by modifying the formula used by Oelz [74]. We then have

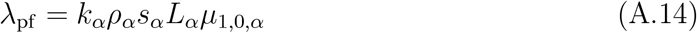

where *k_α_* is the spring stiffness constant of the cross-linker (*α*-actinin), *ρ_α_* is the maximal cross-linker density, *s_α_* is a saturation factor, *L_α_* is the cross-linker length, and *μ*_1,0_ = 1/(*ζ*(1 + *ζ/β*)) is a parameter that incorporates the on-rate, *β*, and off-rate, *ζ*, of the cross-linker, as derived in Milišić and Oelz [38]. Ferrer et al. [75] give *k_α_* = 100pN μm^−1^, and Oelz [74] estimate that *ρ_α_* = 70 μm^−1^ and *s_α_* = 0.05. The length of *α*-actinin is *L_α_* = 36 nm [76]. From Goldmann and Isenberg [77], we obtain an on-rate of *β* = 1 s^−1^, if we assume, as in Oelz, Schmeiser, and Small [78], that the concentration of *α*-actinin is 1 μM. Goldmann and Isenberg [77] also claim that *ζ* = 0.44s^−1^, allowing us to compute *μi,o,a* = 1.5783 s. Thus, *λ*_pf_ = 19.89 pN μm^−1^ s. This is similar in magnitude to the estimate from Ward et al. [49].

#### Myosin Reference Off-Rate, *k*_off_

Stam et al. [59], citing Wang et al. [60] and Kovács et al. [79], state that the reference off-rate *k_o_ff* (0) for non-muscle myosin is 0.35s^−1^ (IIA) and 1.71 s^−1^ (IIB). This parameter therefore depends on the isoform of the myosin, and we adopt the value for myosin-IIA.

#### Actin Turnover Rate, *k*_off,*a*_

In the cell cortex, Saha et al. [43] estimate the timescale for actin filament turnover to be approximately 25 s for *C. elegans* yeast. Based on this, we will use a turnover rate of *k*_off,*a*_ = 0.04 s^−1^ in our simulations.

## B Numerical Simulations and Results

Together with this Supplementary Material, we provide animations and data for the numerical simulations used in this paper. An index of these files appears below.

### B.1 Turnover and Pattern Formation

The following series of plots contains the final network configurations and distance distributions for the ten simulations performed with *T* = 300 s, and both *k*_off,*a*_ = 0s^−1^ and *k*_off,a_ = 0.2 s^−1^.

**Figure B.1:**
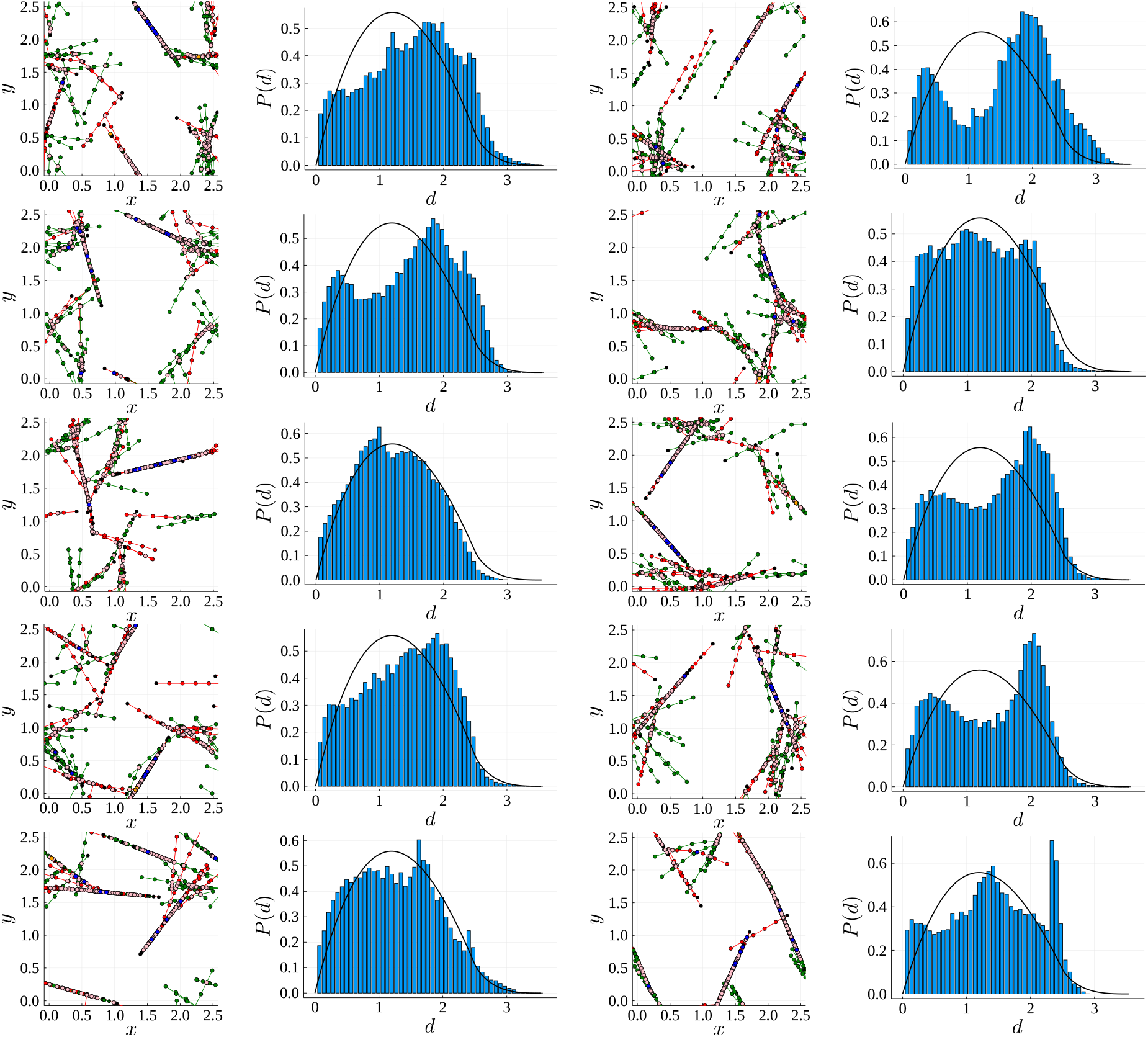
Final network configurations at *t* = 300 s and histograms of the distances between pairs of nodes on different filaments. Results presented for ten simulations with *k*_off,*a*_ = 0s^−1^.

**Figure B.2:**
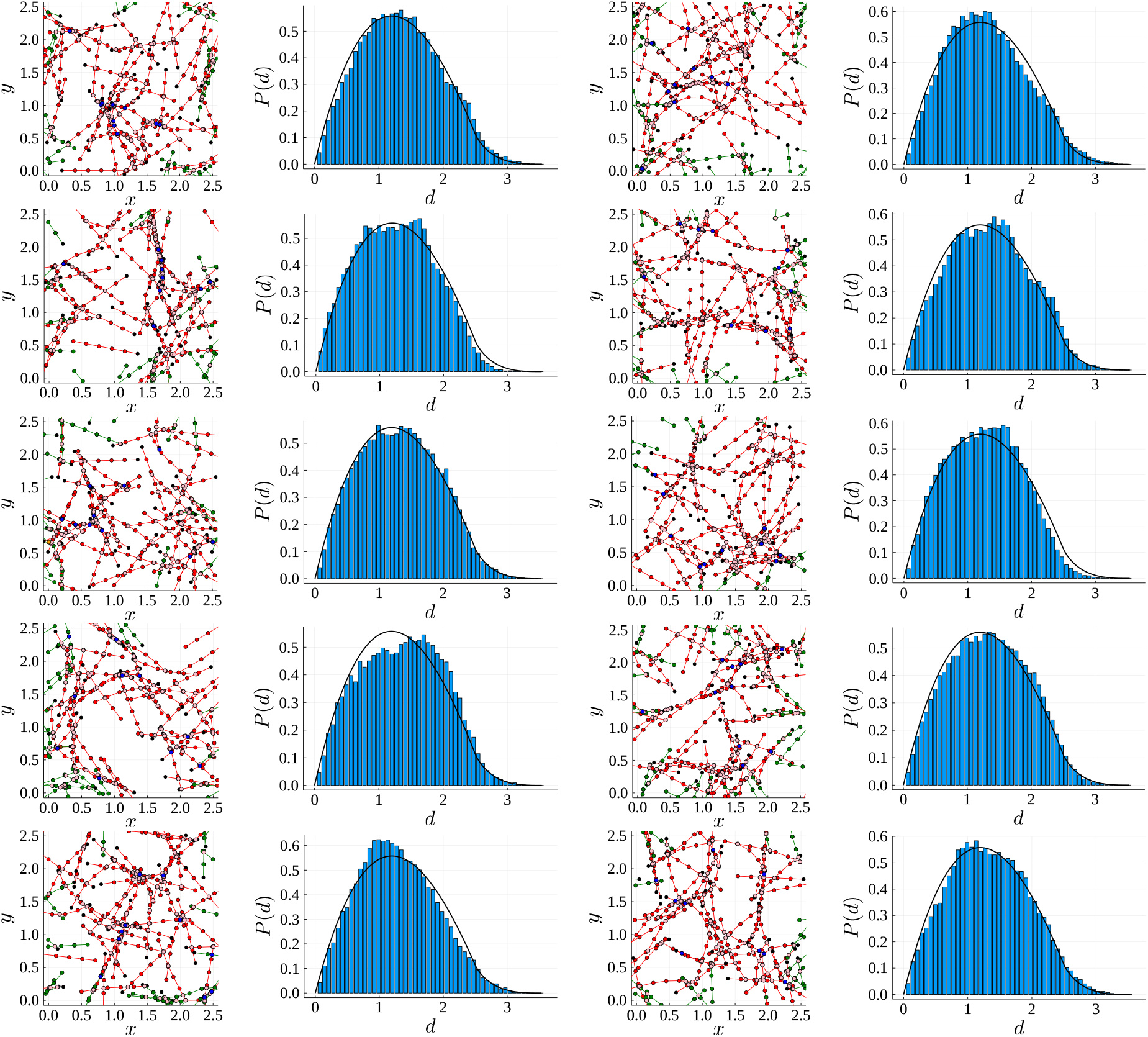
Final network configurations at *t* = 300 s and histograms of the distances between pairs of nodes on different filaments. Results presented for ten simulations with *k*_off,*a*_ 0.2 s^−1^.

## References

[1] M. Gautel, “The sarcomeric cytoskeleton: who picks up the strain?”, Current Opinion in Cell Biology 23 (2011), pp. 39–46, DOI: 10.1016/j.ceb.2010.12.001.

[2] T. D. Pollard, “Mechanics of cytokinesis in eukaryotes”, Current Opinion in Cell Biology 22 (2010), pp. 50–56, DOI: 10.1016/j.ceb.2009.11.010.

[3] K. M. Yamada and M. Sixt, “Mechanisms of 3D cell migration”, Nature Reviews Molecular Cell Biology 20 (2019), pp. 738–752, DOI: 10.1038/s41580-019-0172-9.

[4] K. J. Chalut and E. K. Paluch, “The actin cortex: a bridge between cell shape and function”, Developmental Cell 38 (2016), pp. 571–573, DOI: 10.1016/j.devcel.2016.09.011.

[5] T. D. Pollard, “The value of mechanistic biophysical information for systems-level understanding of complex biological processes such as cytokinesis”, Biophysical Journal 107 (2014), pp. 2499–2507, DOI: 10.1016/j.bpj.2014.10.031.

[6] T. H. Cheffings, N. J. Burroughs, and M. K. Balasubramanian, “Actomyosin ring formation and tension generation in eukaryotic cytokinesis”, Current Biology 26 (2016), R719–R739, DOI: 10.1016/j.cub.2016.06.071.

[7] B. Y. Rubinstein and A. Mogilner, “Myosin clusters of finite size develop contractile stress in 1D random actin arrays”, Biophysical Journal 113 (2017), pp. 937–947, DOI: 10.1016/j.bpj.2017.07.003.

[8] D. Gordon, A. Bernheim-Groswasser, C. Keasar, and O. Farago, “Hierarchical self-organization of cytoskeletal active networks”, Physical Biology 9, 026005 (2012), DOI: 10.1088/1478-3975/9/2/026005.

[9] M. Lenz, “Geometrical origins of contractility in disordered actomyosin networks”, Physical Review X 4, 041002 (2014), DOI: 10.1103/PhysRevX.4.041002.

[10] T. Hiraiwa and G. Salbreux, “Role of turnover in active stress generation in a filament network”, Physical Review Letters 116, 188101 (2016), DOI: 10.1103/PhysRevLett.116.188101.

[11] A. E. Carlsson, “Contractile stress generation by actomyosin gels”, Physical Review E 74, 051912 (2006), DOI: 10.1103/PhysRevE.74.051912.

[12] D. B. Oelz, B. Y. Rubinstein, and A. Mogilner, “A combination of actin treadmilling and cross-linking drives contraction of random actomyosin arrays”, Biophysical Journal 109 (2015), pp. 1818–1829, DOI: 10.1016/j.bpj.2015.09.013.

[13] K. Kruse and F. Jülicher, “Actively contracting bundles of polar filaments”, Physical Review Letters 85 (2000), pp. 1778–1781, DOI: 10.1103/PhysRevLett.85.1778.

[14] M. Lenz, M. L. Gardel, and A. R. Dinner, “Requirements for contractility in disordered cytoskeletal bundles”, New Journal of Physics 14, 033037 (2012), DOI: 10.1088/1367-2630/14/3/033037.

[15] V. Wollrab, J. M. Belmonte, L. Baldauf, M. Leptin, F. Nédeléc, and G. H. Koenderink, “Polarity sorting drives remodeling of actin-myosin networks”, Journal of Cell Science 132, jcs219717 (2019), DOI: 10.1242/jcs.219717.

[16] M. P. Murrell and M. L. Gardel, “F-actin buckling coordinates contractility and severing in biomimetic actomyosin cortex”, Proceedings of the National Academy of Science of the United States of America 109 (2012), pp. 20820–20825, DOI: https://doi.org/10.1073/pnas.1214753109.

[17] J. M. Belmonte, M. Leptin, and F. J. Nédélec, “A theory that predicts behaviors of disordered cytoskeletal networks”, Molecular Systems Biology 13, 941 (2017), DOI: 10.15252/msb.20177796.

[18] T. Kamasaki, M. Osumi, and I. Mabuchi, “Three-dimensional arrangement of F-actin in the contractile ring of fission yeast”, Journal of Cell Biology 178 (2007), pp. 765–771, DOI: 10.1083/jcb.200612018.

[19] E. M. De La Cruz and M. L. Gardel, “Actin mechanics and fragmentation”, Journal of Biological Chemistry 290 (2015), pp. 17137–17144, DOI: 10.1074/jbc.R115.636472.

[20] K. Kruse and F. Jülicher, “Self-organisation and mechanical properties of active actin filaments”, Physical Review E 67, 051913 (2003), DOI: 10.1103/PhysRevE.67.051913.

[21] A. Zumdieck, K. Kruse, H. Bringmann, A. A. Hyman, and F. Jülicher, “Stress generation and filament turnover during actin ring constriction”, PLoS One 2, e696 (2007), DOI: 10.1371/journal.pone.0000696.

[22] K. Kruse, J. F. Joanny, F. Jülicher, J. Prost, and K. Sekimoto, “Asters, vortices, and rotating spirals in active gels of polar filaments”, Physical Review Letters 92, 078101 (2004), DOI: 10.1103/PhysRevLett.92.078101.

[23] D. B. Oelz and A. Mogilner, “Actomyosin contraction, aggregation and traveling waves in a treadmilling actin array”, Physica D 318–319 (2016), pp. 70–83, DOI: 10.1016/j.physd.2015.10.005.

[24] F. J. Nédélec and D. Foethke, “Collective Langevin dynamics of flexible cytoskeletal fibers”, New Journal of Physics 9, 427 (2007), DOI: 10.1088/1367-2630/9/11/427.

[25] S. L. Freedman, S. Banerjee, G. M. Hocky, and A. R. Dinner, “A versatile framework for simulating the dynamic mechanical structure of cytoskeletal networks”, Biophysical Journal 113 (2017), pp. 448–460, DOI: 10.1016/j.bpj.2017.06.003.

[26] K. Popov, J. E. Komianos, and G. A. Papoian, “MEDYAN: Mechanochemical simulations of contraction and polarity alignment in actomyosin networks”, PLoS Computational Biology 12, e1004877 (2016), DOI: 10.1371/journal.pcbi.1004877.

[27] I. Mendes Pinto, B. Y. Rubinstein, A. Kucharavy, J. R. Unruh, and R. Li, “Actin depolymerization drives actomyosin ring contraction during budding yeast cytokinesis”, Developmental Cell 22 (2012), pp. 1247–1260, DOI: 10.1016/j.devcel.2012.04.015.

[28] M. R. Stachowiak, C. Laplante, H. F. Chin, B. Guirao, E. Karatekin, T. D. Pollard, and B. O’Shaughnessy, “Mechanism of cytokinetic contractile ring constriction in fission yeast”, Developmental Cell 29 (2014), pp. 547–561, DOI: 10.1016/j.devcel.2014.04.021.

[29] T. Kim, “Determinants of contractile forces generated in disorganized actomyosin bundles”, Biomechanics and Modeling in Mechanobiology 14 (2015), pp. 345–355, DOI: 10.1007/s10237-014-0608-2.

[30] P. Chugh, A. G. Clark, M. B. Smith, D. A. D. Cassani, K. Dierkes, A. Ragab, P. P. Roux, G. Charras, G. Salbreux, and E. K. Paluch, “Actin cortex architecture regulates cell surface tension”, Nature Cell Biology 19 (2017), pp. 689–697, DOI: 10.1038/ncb3525.

[31] T. D. Pollard and B. O’Shaughnessy, “Molecular mechanism of cytokinesis”, Annual Review of Biochemistry 88 (2019), pp. 661–689, DOI: 10.1146/annurev-biochem-062917012530.

[32] H. Ennomani, G. Letort, C. Guérin, J. Martiel, W. Cao, F. J. Nédélec, E. M. De La Cruz, M. Théry, and L. Blanchoin, “Architecture and connectivity govern actin network contractility”, Current Biology 26 (2016), pp. 616–626, DOI: 10.1016/j.cub.2015.12.069.

[33] D. A. Head, A. J. Levine, and F. C. MacKintosh, “Distinct regimes of elastic response and deformation modes of cross-linked cytoskeletal and semiflexible polymer networks”, Physical Review E 68, 061907 (2003), DOI: 10.1103/PhysRevE.68.061907.

[34] M. Murrell, P. W. Oakes, M. Lenz, and M. L. Gardel, “Forcing cells into shape: the mechanics of actomyosin contractility”, Nature Reviews Molecular Cell Biology 16 (2015), pp. 486–498, DOI: 10.1038/nrm4012.

[35] T. C. Bidone, W. Jung, D. Maruri, C. Borau, R. D. Kamm, and T. Kim, “Morphological transformation and force generation of active cytoskeletal networks”, PLoS Computational Biology 13, e1005277 (2017), DOI: 10.1371/journal.pcbi.1005277.

[36] M. Lenz, T. Thoresen, M. L. Gardel, and A. R. Dinner, “Contractile units in disordered actomyosin bundles arise from F-actin buckling”, Physical Review Letters 108, 238107 (2012), DOI: 10.1103/PhysRevLett.108.238107.

[37] K. Tawada and K. Sekimoto, “Protein friction exerted by motor enzymes through a weak-binding interaction”, Journal of Theoretical Biology 150 (1991), pp. 193–200, DOI: 10.1016/S0022-5193(05)80331-5.

[38] V. Milišić and D. B. Oelz, “On the asymptotic regime of a model for friction mediated by transient elastic linkages”, Journal de Mathématiques Pures et Appliquées 96 (2011), pp. 484–501, DOI: 10.1016/j.matpur.2011.03.005.

[39] W. M. McFadden, P. M. McCall, M. L. Gardel, and E. M. Munro, “Filament turnover tunes both force generation and dissipation to control long-range flows in a model actomyosin cortex”, PLoS Computational Biology 13, e1005811 (2017), DOI: 10.1371/journal.pcbi.1005811.

[40] N. L. Dasanayake, P. J. Michalski, and A. E. Carlsson, “General mechanism of actomyosin contractility”, Physical Review Letters 107, 118101 (2011), DOI: 10.1103/PhysRevLett.107.118101.

[41] G. I. Bell, “Models for the specific adhesion of cells to cells”, Science 200 (1978), pp. 618–627, DOI: 10.1126/science.347575.

[42] G. H. Koenderink and E. K. Paluch, “Architecture shapes contractility in actomyosin networks”, Current Opinion in Cell Biology 50 (2018), pp. 79–85, DOI: 10.1016/j.ceb.2018.01.015.

[43] A. Saha, M. Nishikawa, M. Behrndt, C. Heisenberg, F. Jülicher, and S. W. Grill, “Determining physical properties of the cell cortex”, Biophysical Journal 110 (2016), pp. 1421–1429, DOI: 10.1016/j.bpj.2016.02.013.

[44] G. Salbreux, G. Charras, and E. Paluch, “Actin cortex mechanics and cellular morphogenesis”, Trends in Cell Biology 22 (2012), pp. 536–545, DOI: 10.1016/j.tcb.2012.07.001.

[45] T. Kim, M. L. Gardel, and E. Munro, “Determinants of fluidlike behaviour and effective viscosity in cross-linked actin networks”, Biophysical Journal 106 (2014), pp. 526–534, DOI: 10.1016/j.bpj.2013.12.031.

[46] M. Mak, M. H. Zaman, R. D. Kamm, and T. Kim, “Interplay of active processes modulates tension and drives phase transition in self-renewing, motor-driven cytoskeletal networks”, Nature Communications 7, 10323 (2016), DOI: 10.1038/ncomms10323.

[47] V. Bormuth, V. Varga, J. Howard, and E. Schäffer, “Protein friction limits diffusive and directed movements of kinesin motors in microtubules”, Science 325 (2009), pp. 870–873, DOI: 10.1126/science.1174923.

[48] N. Yoshinaga and P. Marcq, “Contraction of cross-linked actomyosin bundles”, Physical Biology 9 (2012), DOI: 10.1088/1478-3975/9/4/046004.

[49] A. Ward, F. Hilitski, W. Schwenger, D. Welch, A. W. C. Lau, V. Vitelli, L. Mahadevan, and Z. Dogic, “Solid friction between soft filaments”, Nature Materials 14 (2015), pp. 583–588, DOI: 10.1038/nmat4222.

[50] S. Lin, X. Han, G. C. P. Tsui, D. Hui, and L. Gu, “Active stiffening of F-actin network dominated by structural transition of actin filaments into bundles”, Composites Part B: Engineering 116 (2017), pp. 377–381, DOI: 10.1016/j.compositesb.2016.10.079.

[51] P. K. Mogensen and A. N. Risbeth, “Optim: A mathematical optimization package for Julia”, Journal of Open Source Software 3, 615 (2018), DOI: 10.21105/joss.00615.

[52] F. Gittes, B. Mickey, J. Nettleton, and J. Howard, “Flexural rigidity of microtubules and actin filaments measured from thermal fluctuations in shape”, Journal of Cell Biology 120 (1993), pp. 923–934, DOI: 10.1083/jcb.120.4.923.

[53] H. C. Berg, Random Walks in Biology, Princeton University Press, 1993.

[54] D. B. Oelz, U. del Castillo, V. I. Gelfand, and A. Mogilner, “Microtubule dynamics, kinesin-1 sliding, and dynein action drive growth of cell processes”, Biophysical Journal 115 (2018), pp. 1614–1624, DOI: 10.1016/j.bpj.2018.08.046.

[55] T. Thoresen, M. Lenz, and M. L. Gardel, “Reconstitution of contractile actomyosin bundles”, Biophysical Journal 100 (2011), pp. 2698–2705, DOI: 10.1016/j.bpj.2011.04.031.

[56] E. M. Reichl, Y. Ren, M. K. Morphew, M. Delannoy, J. C. Effler, K. D. Girard, S. Divi, P. A. Iglesias, S. C. Kuo, and D. N. Robinson, “Interactions between myosin and actin crosslinkers control cytokinesis contractility dynamics and mechanics”, Current Biology 18 (2008), pp. 471–480, DOI: 10.1016/j.cub.2008.02.056.

[57] J. Wu and T. D. Pollard, “Counting cytokinesis proteins globally and locally in fission yeast”, Science 310 (2005), pp. 310–314, DOI: 10.1126/science.1113230.

[58] K. Murthy and P. Wadsworth, “Myosin-II-dependent localization and dynamics of F-actin during cytokinesis”, Current Biology 15 (2005), pp. 724–731, DOI: 10.1016/j.cub.2005.02.055.

[59] S. Stam, J. Alberts, M. L. Gardel, and E. Munro, “Isoforms confer characteristic force generation and mechanosensation by Myosin II filaments”, Biophysical Journal 108 (2015), pp. 1997–2006, DOI: 10.1016/j.bpj.2015.03.030.

[60] F. Wang, M. Kovács, A. Hu, J. Limouze, E. V. Harvey, and J. R. Sellers, “Kinetic mechanism of non-muscle myosin IIB”, Journal of Biological Chemistry 278 (2003), pp. 27439–27448, DOI: 10.1074/jbc.M302510200.

[61] T. Erdmann and U. S. Schwarz, “Stochastic force generation by small ensembles of myosin II motors”, Physical Review Letters 108, 188101 (2012), DOI: 10.1103/PhysRevLett.108.188101.

[62] C. P. Descovich, D. B. Cortes, S. Ryan, J. Nash, L. Zhang, P. S. Maddox, F. J. Nédélec, and A. S. Maddox, “Cross-linkers both drive and brake cytoskeletal remodeling and furrowing in cytokinesis”, Molecular Biology of the Cell 29 (2018), pp. 622–631, DOI: 10.1091/mbc.E17-06-0392.

[63] S. L. Freedman, G. M. Hocky, S. Banerjee, and A. R. Dinner, “Nonequilibrium phase diagrams for actomyosin networks”, Soft Matter 14 (2018), pp. 7740–7747, DOI: 10.1039/c8sm00741a.

[64] F. J. Nédélec, T. Surrey, A. C. Maggs, and S. Leibler, “Self-organisation of microtubules and motors”, Nature 389 (1997), pp. 305–308, DOI: 10.1038/38532.

[65] M. R. Stachowiak, P. M. McCall, T. Thoresen, H. E. Balcioglu, L. Kasiewicz, M. L. Gardel, and B. O’Shaughnessy, “Self-organization of myosin II in reconstituted actomyosin bundles”, Biophysical Journal 103 (2012), pp. 1265–1274, DOI: 10.1016/j.bpj.2012.08.028.

[66] E. W. Weisstein, Square Line Picking, MathWorld — a Wolfram Web Resource, URL: https://mathworld.wolfram.com/SquareLinePicking.html.

[67] A. C. Martin, M. Kaschube, and E. F. Wieschaus, “Pulsed contractions of an actin–myosin network drive apical constriction”, Nature 457 (2009), pp. 495–499, DOI: 10.1038/nature07522.

[68] L. He, X. Wang, H. L. Tang, and D. J. Montell, “Tissue elongation requires oscillating contractions of a basal actomyosin network”, Nature Cell Biology 12 (2010), pp. 1133–1142, DOI: 10.1038/ncb2124.

[69] Q. Yu, J. Li, M. P. Murrell, and T. Kim, “Balance between force generation and relaxation leads to pulsed contraction of actomyosin networks”, Biophysical Journal 115 (2018), pp. 2003–2013, DOI: https://doi.org/10.1016/j.bpj.2018.10.008.

[70] S. Pellegrin and H. Mellor, “Actin stress fibres”, Journal of Cell Science 120 (2007), pp. 3491–3499, DOI: 10.1242/jcs.018473.

[71] A. Kishino and T. Yanagida, “Force measurements by micromanipulation of a single actin filament by glass needles”, Nature 334 (1988), pp. 74–76, DOI: 10.1038/334074a0.

[72] I. Y. Wong, M. L. Gardel, D. R. Reichman, E. R. Weeks, M. T. Valentine, A. R. Bausch, and D. A. Weitz, “Anomalous diffusion probes microstructure dynamics of entangled F-actin networks”, Physical Review Letters 92, 178101 (2004), DOI: 10.1103/PhysRevLett.92.178101.

[73] I. Obodovskiy, “Chapter 34 — Basics of Biology”, Radiation, ed. by I. Obodovskiy, Elsevier, 2019, pp. 429–445, ISBN: 978-0-444-63979-0, DOI: https://doi.org/10.1016/B978-0-444-63979-0.00034-3.

[74] D. B. Oelz, “A viscous two-phase model for contractile actomyosin bundles”, Journal of Mathematical Biology 68 (2014), pp. 1653–1676, DOI: 10.1007/s00285-013-0682-6.

[75] J. M. Ferrer, H. Lee, J. Chen, B. Pelz, F. Nakamura, R. D. Kamm, and M. J. Lang, “Measuring molecular rupture forces between single actin filaments and actin-binding proteins”, Proceedings of the National Academy of Science of the United States of America 105 (2008), pp. 9221–9226, DOI: 10.1073/pnas.0706124105.

[76] D. H. Wachsstock, W. H. Schwarz, and T. D. Pollard, “Affinity of *α*-actinin for actin determines the structure and mechanical properties of actin filament gels.”, Biophysical Journal 65 (1993), pp. 205–214, DOI: 10.1016/S0006-3495(93)81059-2.

[77] W. H. Goldmann and G. Isenberg, “Analysis of filamin and *α*-actinin binding to actin by the stopped flow method”, Federation of European Biochemical Societies Letters 336 (1993), pp. 408–410, DOI: 10.1016/0014-5793(93)80847-N.

[78] D. B. Oelz, C. Schmeiser, and J. V. Small, “Modeling of the actin-cytoskeleton in symmetric lamellipodial fragments”, Cell Adhesion & Migration 2 (2008), pp. 117–126, DOI: 10.4161/cam.2.2.6373.

[79] M. Kovács, F. Wang, A. Hu, Y. Zhang, and J. R. Sellers, “Functional divergence of human cytoplasmic myosin II”, Journal of Biological Chemistry 278 (2003), pp. 38132–38140, DOI: 10.1074/jbc.M305453200.

